# Sequence design for three-dimensional genome folding using Akita Semifreddo

**DOI:** 10.64898/2026.04.28.721368

**Authors:** Paulina N. Smaruj, David R. Kelley, Geoffrey Fudenberg

**Affiliations:** Department of Quantitative and Computational Biology, University of Southern California, Los Angeles, CA, USA; Calico Life Sciences LLC, South San Francisco, CA, USA

## Abstract

Mammalian genomes display complex three-dimensional organization which is crucial for downstream processes like gene regulation. Local features of genome organization are largely driven by loop extrusion and manifest as boundaries, dots, and flames in genome contact maps. Still, the rational design of DNA sequences that produce desired folding patterns has not been demonstrated. Here, we present Akita Semifreddo, a framework that enables the rational *in silico* design of DNA sequences with programmable 3D folding outcomes. This combines a computationally efficient “half-frozen” version of the AkitaV2 genome folding model with the Ledidi sequence optimizer. We systematically demonstrate that this approach spans the full repertoire of known local folding features. We show that ∼2 kb synthetic sequences can be designed to induce boundaries, dots, and flames at desired strengths, with CTCF motif configurations consistent with their known mechanistic bases. We further demonstrate that weak boundaries can be designed through transcription-associated sequence features alone, without introducing CTCF motifs, and that strong boundaries can be suppressed by introducing SINE B2 retroelement-like sequences. Collectively, these results reveal a many-to-one relationship between DNA sequence and folding outcomes and uncover the biological basis of sequence features leveraged by our model. In short, Akita Semifreddo provides a platform for dissecting the sequence grammar of three-dimensional chromatin architecture and engineering synthetic regulatory landscapes.

## Introduction

Genomes are densely packed with regulatory elements, which often communicate over large genomic distances to control gene expression [1–4]. Gene regulatory elements, including promoters and enhancers, can be separated by hundreds of kilobases or more, yet are brought into proximity by the genome folding machinery [5,6]. The same machinery creates distinct folding patterns — including: boundaries (between neighboring domains), flames (typically appearing at domain edges, also known as stripes), and dots (punctae also referred to as loops) [7]. Positions of these patterns are dictated by DNA sequence features and are thought to guide regulatory communication across the genome [8]. A central challenge for advancing genome biology is to translate our growing understanding of genome folding and regulatory element function into the ability to precisely design DNA sequences *in silico* that achieve desired gene expression outcomes [9–11].

An emerging avenue for *in silico* DNA design makes use of deep neural networks that map DNA sequence to epigenomic profiles [2]. Such approaches fall into three general categories: (i) heuristic strategies [12–15], (ii) generative neural networks [13,16–23], and (iii) optimization-based methods [15,24–26], each with distinct strengths and weaknesses [27]. Heuristic strategies provide intuitive, stepwise control over sequence properties and mechanistic interpretability, yet may not explore the full sequence space. Generative neural networks can efficiently explore vast regions of sequence space, yet are typically constrained to short sequence lengths and can be difficult to train due to architectural complexity and instability (as in GANs) or substantial memory demands (as in diffusion models) [27,28]. Optimization-based methods offer precise control over design objectives though traditional strategies emphasizing local improvements may fail to identify good designs that require expansive exploration of sequence space [27]. Despite the availability of neural networks capable of accurately predicting complex genome folding patterns from DNA sequence inputs, reviewed in [29] and including even more recent models [30,31], designing DNA sequences that specify genome folding remains largely unexplored.

A promising approach for tackling the complexities of genome folding design *in silico* is the Ledidi [26] optimization approach, which generates interpretable designs within a flexible PyTorch framework. The method formulates sequence design as an optimization problem that balances two objectives: preserving the original input (input loss) while steering the model prediction toward a desired output (output loss). The balance between these two terms is controlled by a scaling parameter (λ). Ledidi uses a straight-through Gumbel-Softmax reparameterization, enabling gradient-based updates over discrete nucleotide categories [32], with a temperature parameter (τ) controlling the sharpness of this distribution. While Ledidi was originally applied to 1D genomic profiles, such as transcription factor binding or accessibility tracks, applying it to genome folding presents unique challenges. Contact map predictors, like AkitaV2, operate on much longer input sequences and generate 2D structural features — boundaries, dots, and flames — rather than 1D peaks.

Engineering sequences to produce specific genome-folding patterns presents challenges beyond those encountered for short regulatory elements. First, genome folding arises from interactions spanning hundreds of kilobases to megabases, requiring models that operate over substantially larger sequence contexts. Second, experimental characterization of genome folding is constrained by the relatively coarser resolution of Hi-C and Micro-C data relative to other 1D epigenomic profiles. This limits current predictive models, which cannot achieve base-pair precision, making it difficult to attribute specific sequence edits to changes in the resulting contact maps. Third, the design objective shifts from specifying 1D regulatory profiles to defining 2D contact maps, adding complexity target representation and loss quantification. Finally, the sequence determinants of higher-order chromatin architecture — including motif syntax, cooperative CTCF interactions, and extrusion barriers — remain incompletely understood, which both limits heuristic sequence design approaches and motivates the need for data-driven, model-guided optimization frameworks.

Here, we present Akita Semifreddo (Akita^SF^), a partially frozen version of AkitaV2 that combines seamlessly with the Ledidi sequence optimizer to enable DNA design. We reimplemented AkitaV2 in PyTorch [33] and systematically explored optimization strategies to enable the design of contact map features with Ledidi. Akita^SF^ enables the design of specific sequence segments while accounting for the full >1 Mb genomic context, greatly improving the efficiency of iterative optimization with Ledidi. Our approach enables the design of synthetic DNA sequences that induce key features of local genome folding. Together, these advances provide a framework for rationally designing genomic sequences to manipulate three-dimensional chromatin architecture, opening new avenues for genome engineering, gene regulation, and synthetic biology.

## Results

### Transferring AkitaV2 to the PyTorch framework

To interface with the rich PyTorch ecosystem, including Ledidi, we reimplemented the AkitaV2 architecture in PyTorch. We maintained the exact architecture of the TensorFlow original [33], except setting the output dimension to one and training independent models for each cell type. We directly assigned pretrained TensorFlow weights to the corresponding PyTorch layers for all eight cross-validation folds and the additional cell types used in AkitaV2. The ported PyTorch implementation showed a minor performance reduction relative to the TensorFlow model (**Additional file 1: Table S1**), likely due to subtle numerical and implementation differences between frameworks. To mitigate this, we fine-tuned each PyTorch model from the transferred TensorFlow weights for the corresponding cell type. The validation loss dropped sharply within the first epoch and plateaued, indicating rapid convergence (**Additional file 2: Fig. S1**). Per cell-type fine-tuning restored and often improved performance relative to the TensorFlow AkitaV2 model (**Additional file 1: Table S1**). Benchmarking showed that our fine-tuned PyTorch models achieve better performance than ORCA [34] on human cell types (0.56 vs 0.68 Pearson’s r) and comparable to AlphaGenome [30] on both human and mouse cell types (**Additional file 1: Table S2,** 0.640 vs 0.641 average Pearson’s r for human Hi-C, and 0.75 vs 0.67 for mouse Hi-C). However, our models were slightly less performant than AlphaGenome on human Micro-C datasets (0.68 vs 0.73 Pearson’s r), possibly due to its multimodal training regime, which incorporates chromatin accessibility, RNA-seq, histone modifications, and transcription factor binding data. In this work, we refer to models fine-tuned on mESC data [35] as AkitaPT.

### Ledidi enables complete “rewiring” of genome folding maps

To test Ledidi’s potential for designing genome folding, we paired it with AkitaPT and used the combined framework to perform map-to-map transformations. Specifically, we attempted to redesign input DNA sequences so that their predicted contact frequency maps matched a target map generated from a distinct genomic locus. We permitted edits across the full 1.3 Mb input sequence, providing a proof-of-concept scenario for complete “rewiring” of genome folding.

The transformations were highly successful: optimized maps were visually indistinguishable from their respective targets (**Additional file 2: Fig. S2A**), and accepted sequence edits progressively converged the predicted contact map toward the target (**Additional file 3: Movie S1**). Quantitatively, the majority of optimized maps showed high similarity to their targets (Pearson’s r > 0.9, **Additional file 2: Fig. S2B**), confirming the strong optimization performance of Ledidi with AkitaPT. To assess whether a particular contact map necessitates a unique CTCF pattern, we identified CTCF motif positions in the original, optimized, and target map sequences (**Additional file 2: Fig. S2C**). We then compared the genomic locations and orientations of CTCF motifs. Jaccard indices revealed minimal overlap between optimized and target sequences (**Additional file 2: Fig. S2D**), indicating that multiple distinct CTCF configurations can produce equivalent contact maps — a many-to-one mapping from sequence to folding pattern. In contrast, moderate overlap between the original and optimized sequences suggested that Ledidi tends to preserve the original sequence where possible, minimizing unnecessary edits.

Most optimized sequences contained 100,000s of accepted edits (**Additional file 2: Fig. S2E**), complicating mechanistic interpretation. Moreover, optimizing a full 1.3 Mb sequence imposed a substantial computational and memory burden, requiring thousands of gradient-based updates involving full forward and backward passes through the deep convolutional AkitaPT model. To improve interpretability and efficiency, we next constrained the optimization objective to introduce a single folding feature within an otherwise featureless region, limiting the editable portion of the sequence, and developed methodology to lower the optimization cost.

### Semifreddo enables efficient forward passes during sequence optimization

Ledidi optimization typically requires hundreds to thousands of model evaluations. DNA design in regulatory genomics has traditionally focused on short input sequences — such as enhancers or promoters spanning ∼500 bp — where full forward passes are computationally tractable. In contrast, genome folding features emerge at megabase scales, even when only tens of basepairs are modified. Indeed, current genome-folding models like Akita are memory-intensive because they require large stretches of input DNA (>1Mb). We thus developed Akita^SF^, a “half-frozen” model to interface with Ledidi (**Fig. 1A**). Our strategy exploits the architecture of the Akita convolutional tower, which iteratively pools 1.3 Mb of one-hot encoded DNA into 640 bin-level embeddings, each representing 2048 bp. If edits are restricted to a small subset of bins (e.g., only the central bin), repeatedly re-evaluating the full convolutional tower results in redundant computation. Local input changes only affect bin-level embeddings within the receptive field imposed by pooling (**Fig. 1B**). We thus cached convolutional tower activations and recomputed them only for the bins affected by edits. For predictions within the receptive field radius (±2 bins) to be computed without boundary artifacts, the model requires the DNA sequence of the edited bin plus context spanning the full receptive field on each side (central bin ± 5 bins, totaling 11 bins). In practice, this reduces convolutional tower input from 640 bins to just 11, bypassing most of the memory-intensive operations in the trunk (**Additional file 2: Fig. S3**). Compared to the full AkitaPT model, Akita^SF^ achieves >3× lower peak memory and ∼25% faster runtime, while producing identical predictions (Pearson’s r = 1.0) (**Additional file 1: Table S3**). Akita^SF^ enabled us to leverage Ledidi to efficiently generate DNA designs with desired genome folding patterns without compromising model accuracy.

**Figure 1.**
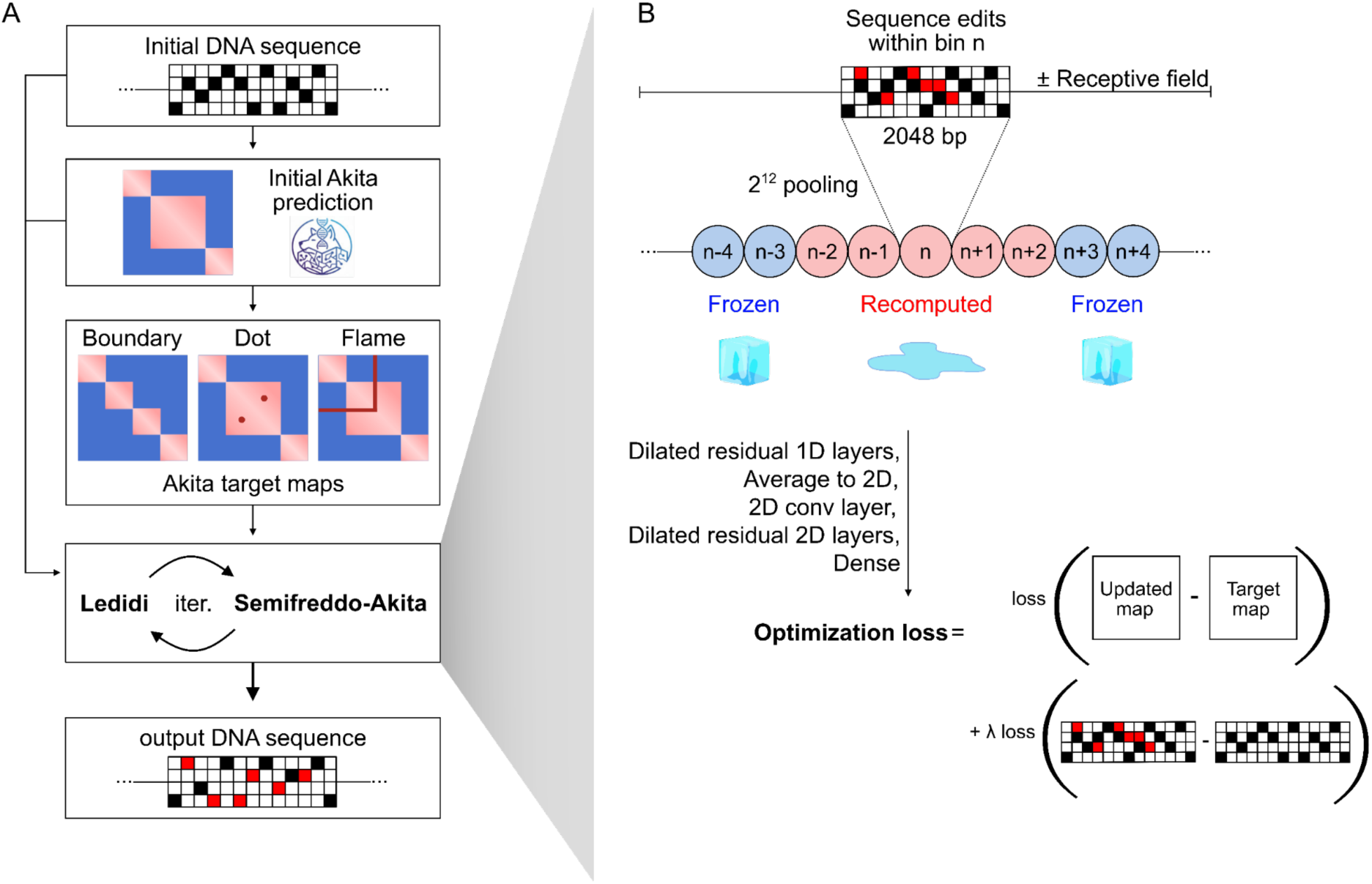
Design of genome folding features using the Akita Semifreddo + Ledidi framework. **A)** The Ledidi optimization algorithm takes as input an initial DNA sequence and a target genome folding map, generated by applying a feature-specific mask to the Akita prediction of the initial sequence. Ledidi iteratively queries the model and accepts only those sequence edits that bring the model’s prediction closer to the desired target. **B)** Akita Semifreddo (Akita^SF^), a “half-frozen” model, recomputes only a subset of activations: edits are restricted to bin *n*, and due to the model’s receptive field (∼5 bins, each 2,048 bp), only bins *n-2* to *n+2* are affected, while the majority of activations at the end of the convolutional tower remain unchanged. By caching convolutional activations and selectively updating outputs for affected bins, the framework efficiently produces edited DNA sequences that achieve the desired genome folding patterns, using less memory and in less time. Finally, the model returns an optimization loss which depends on the difference between the map for the current iteration and target map as well as loss that depends on the number of edits, scaled by an adjustable parameter λ.

### Sequence design for each major feature of local genome folding

To isolate local folding feature design, we used Akita^SF^ to modify structurally “featureless” genomic regions. We took inspiration from Zuin et al. [6], who examined genome folding and transcription within a uniform TAD. After filtering, we obtained 355 featureless genomic regions ranging in size from 205kb to 655kb, which we confirmed through visual inspection (**Methods**, **Additional file 2: Fig. S4A,B**). We systematically explored the design of three characterized genome folding patterns (**Fig. 2**). For each feature, we considered how to: (i) specify the target map, (ii) define the region of the target map used for the output loss function, and (iii) choose optimization hyperparameters controlling the edit count and success rate. We centered output maps to align designed features with the center of the flat regions (**Additional file 2: Fig. S4A**). This approach yielded high success rates for all three features (**Additional file 1: Table S4-6**), with representative examples shown in **Figure 2**.

**Figure 2.**
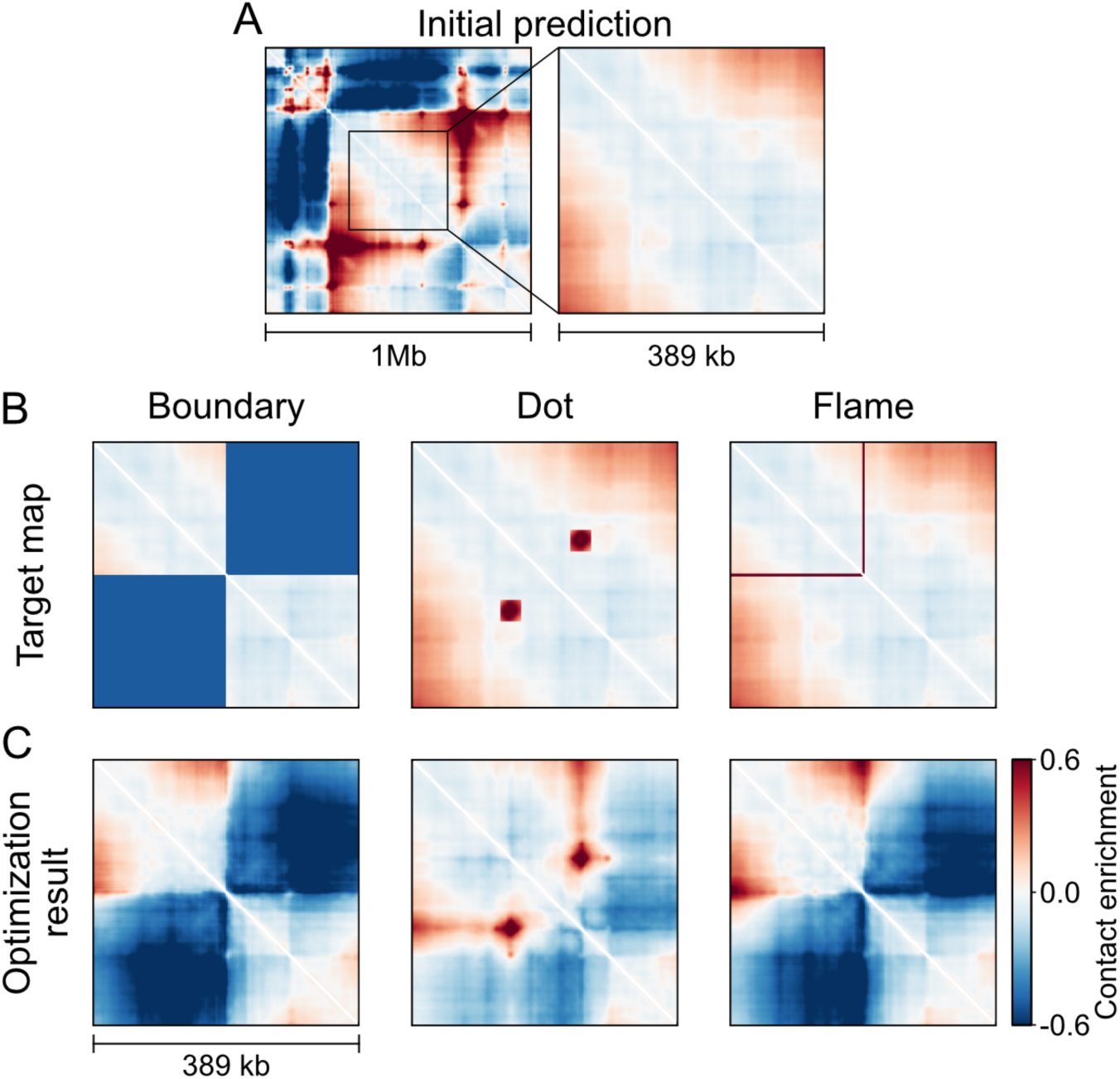
Design of three genome folding features: boundaries, dots, and flames. **A)** Designs were initialized from genomic mouse DNA sequences predicted to have a flat central region (>200 kb). The left panel shows an example genomic contact map, with a magnified view of the flat region shown at right. **B)** From left to right, columns correspond to optimization targets for a boundary, a dot, and a flame. Optimization targets have a feature-specific mask applied to the initial map (boundary and flame: constant target values of −0.5 and 1.0, respectively; dot: data-driven mask). **C)** Resulting optimized contact map for each feature.

We first focused on boundary design, restricting edits to the single central bin of each featureless region. We evaluated target map specification by either adding a pattern mask or rewriting the upper-right quarter of the desired map. We adopted the latter, as computing the output loss over the upper-right quarter proved sufficient to guide boundary formation.

Hyperparameter analysis revealed that increasing the input loss scaling (λ) reduced map alteration, post-optimization CTCF motifs, and success rate. As λ increased from 0.01 to 200, the average number of central bin edits decreased from 792 to 32. An intermediate scaling (λ=125) balanced these factors, reducing edits to ∼51 while maintaining a high success rate (**Additional file 1: Table S4**). At fixed λ, sequences for which optimization failed tended to exhibit lower pre-existing CTCF information content, as measured by positive motif scores aggregated along the optimized sequence (**Additional file 2: Fig. S5A**). When rerun, more than half of the failed optimizations completed successfully, indicating that input loss scaling could be relaxed for sequences with fewer initial CTCF-like subsequences. When the same set of optimizations was executed across 10 independent runs, the number of successful optimizations correlated with pre-existing CTCF information content (**Additional file 2: Fig. S5B**), suggesting that sequences with lower initial CTCF-like content may require multiple rounds of stochastic optimization to achieve success when a higher λ is used. Tuning other Ledidi design parameters (ε and τ) was less effective: lowering ε did not significantly reduce edit number (**Additional file 1: Table S7**), while tuning τ to lower edits yielded optimizations with near-zero average boundary scores (mean = −0.07). Among successful strong boundary designs (n=323, target boundary score = −0.5), every resulting sequence was unique. Designed sequences had ∼3 CTCF motifs on average, and each motif (including ±15 bp flanks) was distinct.

We next explored the design of dots and flames. For dots, edits were allowed in two bins corresponding to dot anchors; for flames, in a single bin. In all cases, rewriting a portion of the map to include the desired feature, combined with a local output loss restricted to that region, improved design success. We optimized λ individually for each feature to achieve approximately 50 edits per bin on average (**Additional file 1: Table S5,6**).

Finally, we validated our designs *in silico* by assessing whether AlphaGenome also predicted the emergence of synthetic features generated by Akita^SF^. For each feature, we assessed concordance both visually and quantitatively. All well-characterized genome folding features — boundaries, dots, and flames — validated with high concordance (**Additional file 2: Fig. S6**). For dot designs, where the localized dot score showed lower cross-model agreement, contact map root mean squared difference (RMSD) revealed substantially higher concordance (**Additional file 2: Fig. S6G**), suggesting that the discrepancy likely reflects slight differences in the predicted signal strength of the surrounding regions between the two models, which become compounded when measuring change over a small localized area rather than the broader map. In contrast with these well-characterized and prevalent features, designs requesting certain less prevalent features, did not validate with AlphaGenome. This included fountains (also termed jets) identified in early zebrafish embryos [36], in thymocytes and resting B-cells [37] and upon WAPL depletion [38] (**Additional file 2: Fig. S7**).

### Sequence features of designed boundaries scale with their targeted strength

In Akita^SF^, a key component of feature design is their requested strength. We thus assessed how designed boundary strength scales with requested boundary strength in target maps. We determined a representative range for requested boundary strength by quantifying the strength of ∼4000 natural genomic boundaries (**Methods**). We selected four target values spanning this natural range and successfully designed boundaries for each. Stronger target boundaries produced correspondingly stronger optimized boundaries (**Fig. 3A**) and were associated with larger CTCF clusters (**Fig. 3B**), consistent with CTCF’s established role in boundary formation. Tracking the optimization trajectory revealed that edits accumulate preferentially around CTCF motif sites, with accepted edits converging on positions that ultimately harbor CTCF motifs in the final sequence (**Fig. 3C**). Analysis of CTCF motif orientations revealed that both left- and right-oriented sites contribute to designed boundaries (**Additional file 2: Fig. S8A**).

**Figure 3.**
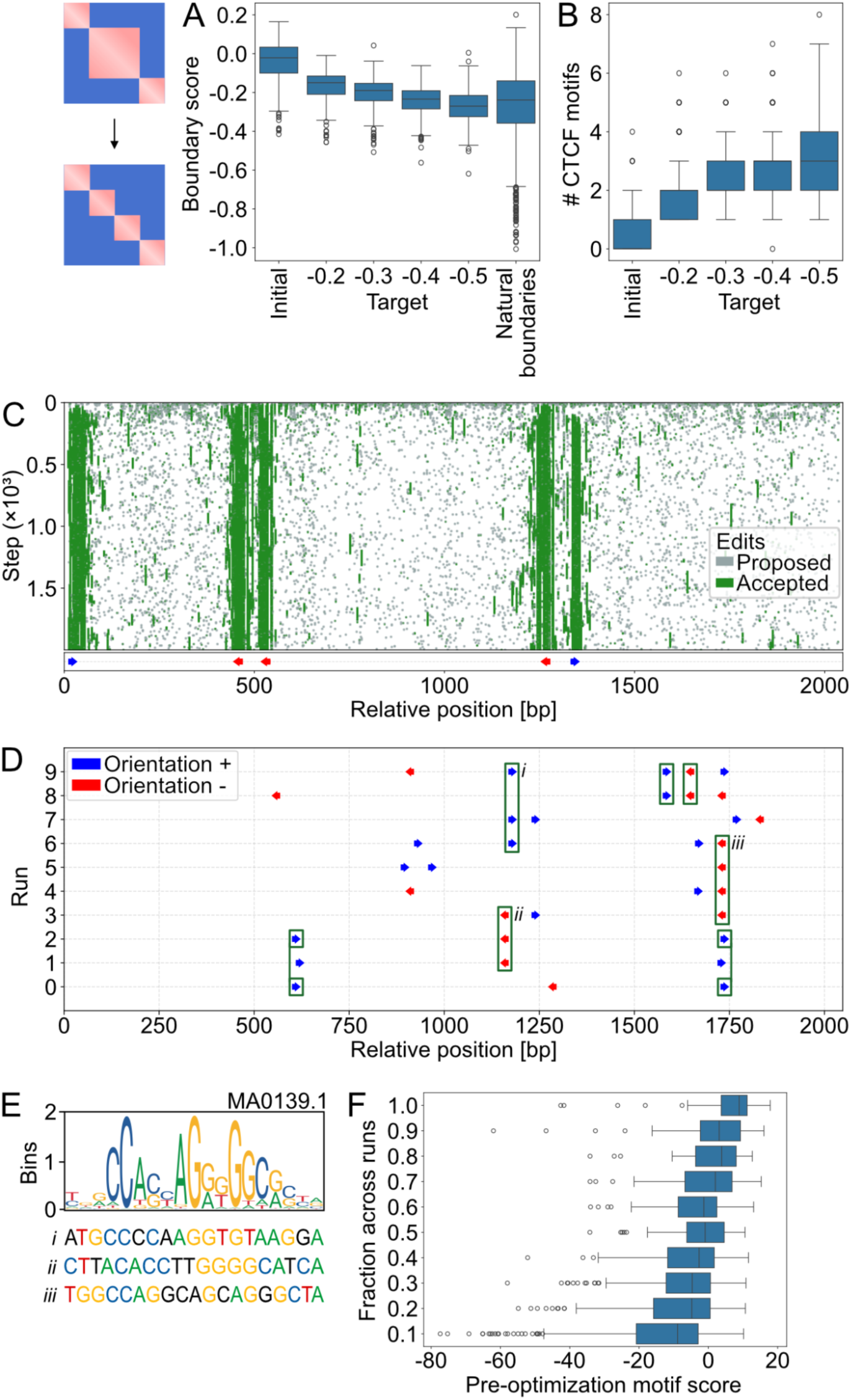
Sequence features of designed boundaries. **A)** Boundary scores, defined as the average predicted signal in the upper-right quarter of the contact map (lower values indicate stronger boundaries), shown for initial mouse genomic DNA sequences (optimization starting points), four target boundary scores (−0.2, −0.3, −0.4, −0.5), and natural boundaries (estimated from >4,000 mouse boundaries). **B)** Number of CTCF motifs detected in the initial sequences and in the designed sequences for the four target boundary strengths. **C)** Edit history and final CTCF motif positions for a single boundary optimization run. The y-axis corresponds to the optimization iteration, and the x-axis to the position within the edited 2,048 bp sequence. Each grey dot indicates an edit proposed at a given iteration and position; each green dot indicates an edit that was accepted. Note that accepted edits can be subsequently reversed. Below the edit history, CTCF motif positions and orientations detected in the final designed sequence are shown as arrows (blue, forward strand; red, reverse strand). Edits accumulate to yield the final set of CTCF motif sites. **D)** CTCF motif positions detected in ten independent optimizations for a strong boundary (target = −0.5) within the same genomic region. Each row represents an independent run, covering the edited central bin. Detected CTCF motifs are indicated by arrows (blue, forward; red, reverse). Green rectangles highlight repeated motif positions and orientations across runs. **E)** CTCF logo (MA0139.1, JASPAR [63]) and sequences prior to optimization at the three most frequent CTCF locations from D. Nucleotides matching the first or second most preferred nucleotide in the logo are color-coded. **F)** Quantification of CTCF motif usage versus pre-optimization CTCF motif similarity. Sites that became CTCF motifs in at least one run were binned by their usage frequency and the distribution of CTCF motif scores for each frequency range is displayed as a boxplot. Less frequently used motifs had systematically lower motif scores before optimization.

We next tested whether boundaries with extreme strength could be designed. Optimization toward extreme targets plateaued both for boundary strength and the number of new CTCF motifs (**Additional file 2: Fig. S8B,C**). This saturation likely reflects optimization dynamics rather than an inherent limit of the model, as we previously observed increasing boundary strength with additional co-oriented motifs at user-defined spacings [33]. Designing densely packed, co-oriented CTCF arrays would require the optimizer to reposition and reorient existing motifs. Such “recombination”-like moves are not accessible to the current gradient-based approach, limiting its ability to reach such highly ordered motif configurations.

### Boundary designs use a flexible set of motif positions influenced by existing sequence variation

Repeated boundary optimizations initialized from the same sequence produced similar boundary scores (**Additional file 2: Fig. S8D**). This prompted us to ask whether optimization returns identical or diverse sequences. To address this, we compared CTCF placement across independent runs, considering motifs as overlapping if both position and orientation matched (**Fig. 3D**). We observed that multiple distinct CTCF configurations can achieve equivalent boundary strength from the same initial sequence. Across optimizations, certain motif locations and orientations appeared preferentially, whereas others were rare and appeared only once among 10 independent optimizations.

The most frequently used CTCF motif positions in optimized sequences displayed some similarity to the canonical CTCF motif prior to optimization (**Fig. 3E**). To quantify this, we computed a CTCF motif score, as in [39,40], for sub-sequences that became CTCF motifs after optimization (**Methods)**. Higher motif scores, indicating stronger pre-existing CTCF-like sub-sequences, correlated with the frequency of CTCF motif incorporation during optimization (**Fig. 3F**). Our finding for genome folding designs aligns with previous observations for 1D ChIP profile design, where Ledidi efficiently activates sequences that are a few edits away from CTCF motifs [26].

To quantify motif positioning diversity across optimization runs for a given initial sequence, we calculated the Jaccard index of CTCF motif usage overlap between every pair of successful designs. Genomic loci displayed a wide range of Jaccard indices (**Additional file 2: Fig. S8E**). Loci with higher pre-existing CTCF information content, as measured by positive motif scores aggregated along the optimized sequence, tended to reuse specific motif positions, resulting in higher average Jaccard indices (**Additional file 2: Fig. S8F**). In contrast, motif-poor initial sequences led to designs with more diverse configurations and lower average Jaccard indices. We interpret this to indicate that optimization is guided by an initial sequence’s latent motif content: pre-existing motifs constrain the solution space.

We noticed that the overall diversity of optimized CTCF configurations also depended on the input loss scaling parameter λ. When edit cost was lower, Akita^SF^ explored a wider variety of solutions (**Additional file 2: Fig. S8G**). In summary, while certain CTCF positions are preferentially utilized, the flexibility of motif placement allows multiple distinct configurations to achieve similar boundary strength, with both pre-existing sequence features and optimization parameters governing the reproducibility and diversity of designs.

### Synthetic dots reproduce the convergent CTCF motif pattern

To design dot patterns, we allowed edits in two bins separated by a defined distance and used a data-driven mask to specify the desired dot position and strength (**Fig. 2B**, **Methods**).

Designing dots with ∼50 edits per bin required increasing the input loss scaling parameter λ compared to boundary design (150 vs 125; **Additional file 1: Table S5**). We tested three inter-anchor distances (30, 50, and 70 bins). Dots at shorter distances (30 bins) were more challenging to design, showing lower success rates, and higher total edits (**Additional file 1: Table S9**). This likely reflects the inherent limit on the maximal observed/expected values at short genomic distances (i.e., near the map diagonal) relative to the requested dot strength.

The strength of designed dots was largely independent of inter-anchor distance (**Additional file 2: Fig. S9A**) and overlapped with that of natural dots (**Methods**). Similarly, the average number of detected CTCF motifs in both bins was around two at all tested inter-anchor distances (**Additional file 2: Fig. S9B**). Orientation analysis revealed that the most frequent configuration consisted of at least one “+” motif at the left anchor and at least one “−” motif at the right anchor, reproducing the characteristic convergent (+−) arrangement associated with dot formation [41] (**Additional file 2: Fig. S9C**).

### Unidirectional CTCF orientation agrees with the mechanistic basis of flame formation

To design flame patterns, we first needed to establish a target flame mask based on naturally occurring flames. We quantified the strength of naturally occurring flames predicted by Akita using StripeNN [42], which detected 3,631 flames genome-wide. We then filtered for high-confidence flames based on: low fraction of missing neighbor and overlapping bins, positive stripiness (as recommended by StripeNN), and positive mean signal in AkitaPT predictions. We quantified natural flame strength as the 75th percentile of the predicted signal within the stripe region (**Methods, Additional file 2: Fig. S9D**). We defined our target flame mask as a symmetric 3-bin-wide stripe of contact enrichment with a specified target value (**Fig. 2B**).

To design flames, we allowed edits in the central bin only. Flame designs with ∼50 edits per bin required increasing the input loss scaling parameter λ relative to boundary design (λ = 140 vs. 125; **Additional file 1: Table S6**). We next designed flames with varying requested strengths. Stronger requested flames yielded correspondingly stronger designs (**Additional file 2: Fig. S9D**) and were associated with larger CTCF clusters (**Additional file 2: Fig. S9E**). Orientation analysis of CTCF clusters in designed sequences revealed “−”, “−−”, and “−−−” as the most frequent configurations (**Additional file 2: Fig. S9F**), indicating a bias toward unidirectional motif arrangements. This enrichment of unidirectional CTCF clusters is consistent with the current mechanistic understanding of flame formation: a directional barrier is required to halt cohesin extrusion from one side [7,43].

### Boundary designs without CTCF motifs display signatures of transcription

We next asked whether boundaries could be designed without introducing detectable CTCF motifs. To test this, we included a CTCF motif penalty into the loss, supplementing the typical Ledidi input and output loss terms for Akita^SF^ (**Methods**). With a sufficiently strong penalty, designed sequences contained no detectable CTCF motifs, yet still produced measurable insulation in the predicted contact maps (**Fig. 4A**, **Additional file 2: Fig. S10A**). Overall, we generated 178 CTCF-free sequences exhibiting clear contact depletion, which we analyzed to identify the sequence features that Akita^SF^ leverages. We observed elevated GC content in optimized sequences relative to their original versions (**Additional file 2: Fig. S10B**). However, dinucleotide-preserving shuffling of the optimized sequences abolished the predicted boundaries (**Additional file 2: Fig. S10A**), indicating that the insulation effect depends on specific sequence features beyond GC content alone. Since active promoters and transcription start sites are known to exhibit insulating properties [44–46], we analyzed Puffin-D [47] GRO-cap predictions for both original and optimized sequences. After optimization, sequences were predicted to have increased transcription initiation relative to their original states (**Fig. 4B**), with the median signal rising more than 7-fold, shifting from a negligible level to that of a moderately expressed endogenous promoter (43rd percentile, **Fig. 4B**; **Methods**). We observed similar trends for predicted Puffin-D PRO-cap and FANTOM CAGE signals. These results suggest that weak insulation can emerge through transcription-associated features independent of canonical CTCF motifs, and that deep neural networks can capture these more dispersed regulatory signals. Finally, we validated the CTCF-free insulation predictions using AlphaGenome and observed strong qualitative and quantitative agreement between insulation scores predicted by the two models (Pearson’s r = 0.64, **Additional file 2: Fig. S10C,D**), supporting the biological plausibility of the designs. Ultimately, these findings indicate that Akita^SF^ can leverage sequence features of active transcription to guide boundary design without CTCF motifs.

**Figure 4.**
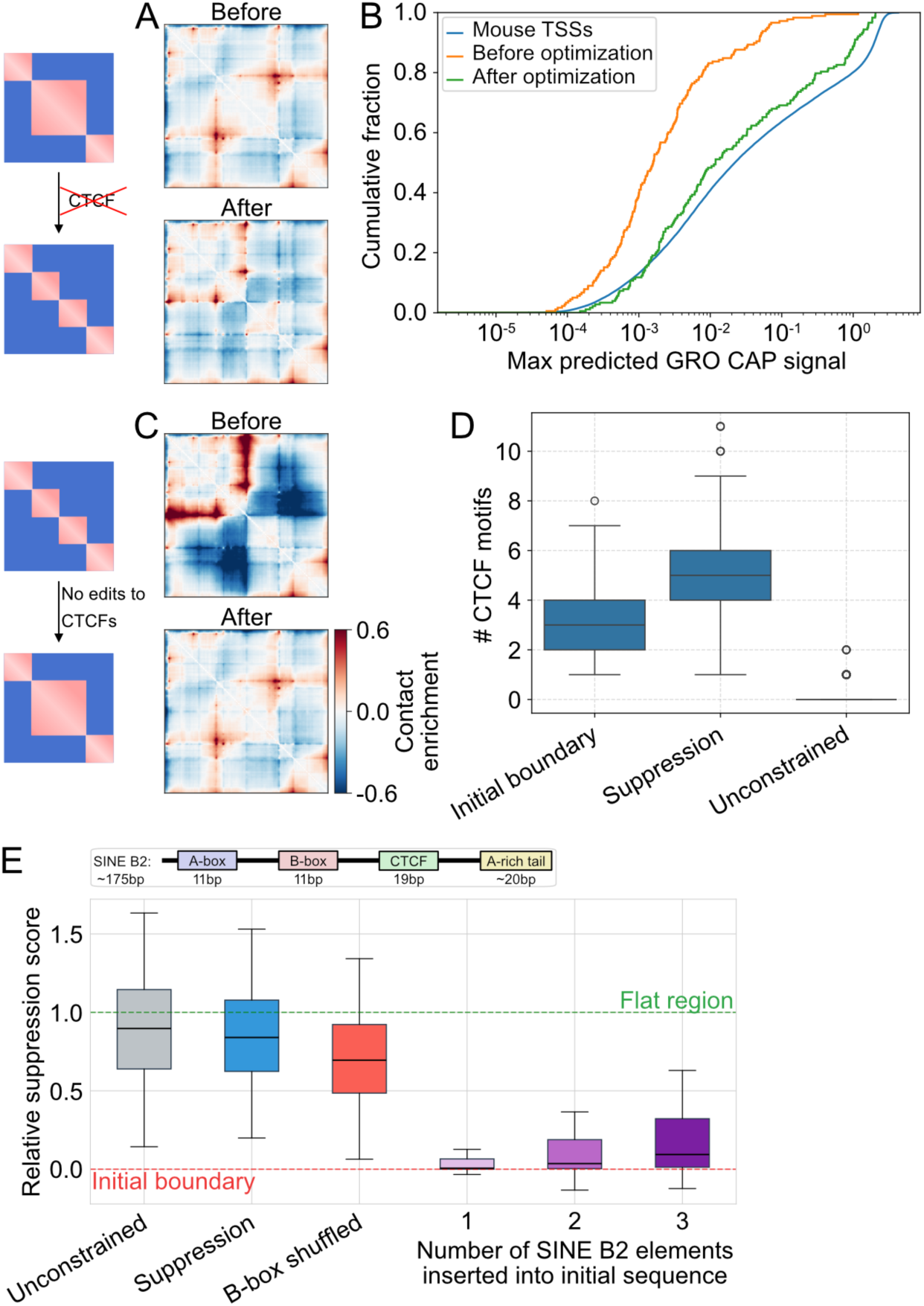
Boundary design without CTCF motifs and boundary suppression. **A)** Predicted contact maps illustrating boundary design without CTCF motifs. *Top:* genomic prediction with a flat central region. *Bottom:* optimized weak boundary (target = −0.2) with no CTCF motifs created. Same color scale as in panel C. **B)** Cumulative fraction of sequences versus maximum Puffin-D predicted GRO-cap signal for 195 designed sequences with detected weak boundaries and no CTCF motifs: before optimization (green), after optimization (orange), and for comparison, maximum GRO-cap predictions for >51,000 mouse TSSs. **C)** Example of boundary suppression without altering pre-existing CTCFs. *Top:* genomic prediction with a synthetic strong boundary. *Bottom:* result of boundary suppression (boundary weakened, pre-existing CTCF motifs unchanged). **D)** Number of CTCF motifs detected in sequences with strong synthetic boundaries before optimization, after locked-motif boundary suppression, and after an unconstrained optimization with CTCF motifs editable as a control. **E)** Relative insulation recovery under different sequence perturbation regimes. *Boxplots from left-to-right:* (i) unconstrained suppression (all motifs editable), (ii) locked-motif suppression, (iii) locked-motif suppression followed by B-box shuffling. B-box shuffling alters boundary strength toward the original baseline. Note that relative suppression can exceed 1.0 in unconstrained scenarios because Akita^SF^ may eliminate endogenous CTCFs present in the original flat genomic regions. *(iv-vi)* relative suppression scores upon insertion of one, two, or three copies of the SINE B2 (B2_Mm2 family) consensus sequence into sequences harboring synthetic boundaries. All endogenous CTCF motifs and ±15bp flanking regions were kept intact during insertion. Boundary strength decreases progressively with insertion number, demonstrating dose-dependent suppression by SINE B2 sequences. *Top:* Schematic of a SINE B2 element highlighting the B-box internal promoter, with segment lengths annotated below.

### Boundary suppression converges on SINE B2-like sequence features

We next asked whether we could identify sequence features that suppress CTCF-mediated boundaries **(Fig. 4C-E**). To test this, we initialized Akita^SF^ with sequences containing strong synthetic boundaries (target = −0.5) and optimized toward their removal — effectively reversing the boundary design process. In an unrestricted optimization where CTCF motifs could be freely edited, the model suppressed boundaries by eliminating CTCF motifs, as expected (**Fig. 4D,E**).

To move beyond this trivial solution, we performed a “locked-motif” optimization, where all CTCF motifs and their flanking regions were excluded from sequence edits (±15 bp, **Methods**). The model still successfully suppressed strong boundaries under these constraints (**Fig. 4C,E**). To our surprise, however, the total number of CTCF motifs increased after optimization (**Fig. 4D**). Boundary suppression was validated using AlphaGenome: optimized sequences showed visually weakened boundaries, and boundary scores predicted by Akita^SF^ and AlphaGenome were modestly correlated (R = 0.45), with insulation consistently decreasing after optimization (**Additional file 2: Fig. S10E,F**).

We next investigated what features the model leveraged to suppress boundaries while concomitantly increasing the number of CTCF motifs. While GC content increased following boundary-suppressing optimization (**Additional file 2: Fig. S10G**), values after editing remained within Akita’s training distribution (**Additional file 2: Fig. S10H**). K-mer analysis revealed enrichment of G and C homopolymers in suppressed sequences. However, replacing these homopolymers with GC dinucleotide repeats — preserving overall GC content — partially restored boundary strength in Akita^SF^, but not in AlphaGenome, suggesting the homopolymer effect is model-specific rather than biologically meaningful.

SINE B2 retroelements are particularly prevalent repetitive elements in the mouse genome and are known to silence their internally embedded CTCF motifs [48–50]. Previous analysis of Akita’s predictions suggested models trained on mouse Hi-C data learn to silence SINE B2-embedded CTCF motifs [51]. We thus investigated whether optimization for boundary suppression generated SINE B2-like features. We found that convolutional filters in Akita activated by SINE B2s were also preferentially activated in boundary-suppressed sequences (**Additional file 2: Fig. S11**). Since the B-box portion of the SINE B2 is involved in silencing by ChAHP *in vivo* [52], we tested whether this contributed to suppression in our designed sequences. We found that model predictions after shuffling B-boxes partially restored boundaries (**Fig. 4E**), further supporting the conclusion that SINE B2 features were leveraged by Akita^SF^ to achieve suppression.

To directly test whether SINE B2 sequences are sufficient to suppress predicted boundaries, we randomly inserted one, two, or three copies of the consensus element, keeping all CTCF motifs and flanking regions intact. SINE B2 sequences weakened boundaries in a dose-dependent manner, though three consensus elements did not recapitulate the full degree of predicted suppression in our designs (**Fig. 4E**) potentially reflecting positional preferences that are not recapitulated with random insertion. Together, these results indicate that Akita^SF^ leverages a biologically-grounded mechanism for boundary suppression, converging on the known ability of SINE B2 elements to silence CTCF-mediated insulation through internally embedded motifs [48,50].

## Discussion

Here, we demonstrate a computationally efficient framework for designing DNA sequences that produce desired genome folding patterns by combining the Akita^SF^ model with the Ledidi optimizer. We developed optimization protocols that successfully produce the major features of local genome folding: boundaries, flames, and dots with desired strengths. Multiple distinct sequences produced equivalent folding patterns, revealing a many-to-one mapping. Boundaries designed without CTCF motifs displayed weak insulation and elevated transcription initiation signals, while boundary suppression experiments converged on SINE B2-like features that silence CTCF-mediated insulation even in the presence of intact CTCF motifs. Each of these sequence designs were computationally validated with AlphaGenome [30], an independently trained model with distinct architecture. In the future, testing designs experimentally, particularly those related to potentially transcription-driven boundaries and boundary suppression, will refine our understanding of sequence-folding relationships. In summary, Akita^SF^ provides a platform for interrogating the sequence grammar of genome folding and engineering synthetic regulatory landscapes.

### Limitations

As for other gradient-based deep-learning design protocols, our framework inherits the limitations of the model used for design as well as the limitations of gradient descent. As limited by AkitaV2, we cannot design features absent from training data or interactions beyond the ∼1.3 Mb prediction window. Attempts to design extreme boundaries plateaued at natural insulation ranges, likely reflecting gradient descent’s inability to efficiently reposition motifs into densely packed, co-oriented CTCF arrays. Additional limitations emerged with rare or chromatin state-driven features: fountain designs failed validation, potentially due to limited training examples and/or diffuse covariates in sequence alone, and cell-type-specific boundaries showed poor fidelity. These results underscore that desired designs require appropriate models and visual validation. Finally, alternative optimization strategies such as reinforcement learning or evolutionary algorithms could overcome gradient descent limitations in designing extreme motif configurations.

## Methods

### Model Finetuning and Benchmarking

#### Akita to Pytorch Transfer

We reimplemented the AkitaV2 architecture in PyTorch by reproducing each layer from the original TensorFlow model. AkitaV2 TensorFlow weights were then transferred directly into the PyTorch implementation, and all tensor shapes, convolutional parameters, and normalization settings were verified to ensure architectural equivalence. Although the transferred model performed similarly to the TensorFlow reference, we observed a slight decrease in accuracy (**Additional file 1: Table S1**). To mitigate this discrepancy, we fine-tuned the PyTorch model starting from the imported TensorFlow weights. Batch-normalization statistics were recalibrated using preciseBN from fvcore.nn.precise_bn [53], which computes population statistics over the full training set. Fine-tuning was performed using a low learning rate (0.001) and the schedule-free AdamW optimizer [54], reflecting that only minimal updates were required to reconcile the two implementations. Because nearly all performance gains occurred within the first epoch (**Additional file 2: Fig. S1**), training was terminated using an early-stopping patience of five epochs. To assess the agreement between the TensorFlow and PyTorch models, we evaluated predictions on held-out data by comparing the vectorized upper-triangular entries of the predicted and target contact-frequency maps. Pearson’s correlation, Spearman’s correlation, and mean squared error (MSE) were computed on concatenated predicted and true vectors. All data preprocessing steps followed the protocol described in the original AkitaV2 publication [33].

#### AlphaGenome and AkitaV2 Mouse Test Set Overlap

We reproduced the AlphaGenome [30] dataset split using the Borzoi [55] sequences_mouse.bed.gz file (https://github.com/calico/borzoi/tree/5c9358222b5026abb733ed5fb84f3f6c77239b37/data). Following the AlphaGenome protocol, each 196-kb Borzoi target interval was expanded to a 1 Mb input window centered on the interval midpoint. To maintain strict separation between training and test partitions, we excluded any test interval whose 1 Mb window overlapped a 1 Mb training window within the same fold. This procedure yielded a cleaned test-sequence table for AlphaGenome models 0-3. To identify the subset of AkitaV2 windows that corresponded to AlphaGenome test regions, we intersected this table with the AkitaV2 test sequences using the bioframe [56] overlap function with the “inner” strategy. AkitaV2 windows that overlapped more than one AlphaGenome test window originating from different folds were discarded to preserve fold integrity. For each remaining AkitaV2 window, we retained the AlphaGenome window exhibiting the largest overlap. This filtering resulted in 2,940 matched windows, from which we randomly sampled 500 to evaluate model performance on AkitaV2 prediction windows.

#### AlphaGenome, ORCA and, AkitaV2 Human Test Set Overlap

We applied the same procedure as described for the mouse dataset, using the Borzoi [55] sequences_human.bed.gz file (https://github.com/calico/borzoi/tree/5c9358222b5026abb733ed5fb84f3f6c77239b37/data). Intersecting AlphaGenome and AkitaV2 [33] test windows produced 2,753 overlapping regions. To focus on sequences held out for ORCA [34] evaluation, we further restricted this set to sequences located on chromosomes 9 and 10, yielding 352 windows. These sequences were subsequently used to assess model performance on AkitaV2 prediction windows.

For ORCA evaluation, each 1,310,720 bp AkitaV2 test window was extended by 344,640 bp on each side to match ORCA’s required 2,000,000 bp input. ORCA produces a 500×500 contact map at 4 kb resolution; to enable direct comparison with Hi-C targets at 2,048 bp resolution, the central 262×262 bins of each predicted map were cropped and subsequently upsampled to 512×512 using bilinear interpolation (zoom_array from cooltools [57]), matching the resolution and dimensions of the AkitaV2 and ORCA predicted matrices.

### Genomic Loci Selection for *in silico* DNA design

To identify relatively “featureless” candidate regions for *in silico* design, we applied a multi-step filtering procedure (**Additional file 2: Fig. S4A**). For each cross-validation fold, prediction examples with per-sample Pearson’s correlation coefficients exceeding the overall mean were initially selected. For each chosen region, we computed the predicted insulation profile using a diamond-shaped window of 16 bins (∼32.7 kb) and identified the longest continuous segment (≥100 bins, ∼205 kb) exhibiting minimal local variability and located sufficiently far from map edges (>50 bins). Flat regions were detected using a sliding-window approach, which flags segments where local variability remains below a defined threshold while preserving maximal region length and avoiding edge-proximal segments. Each candidate region was further verified through visual inspection of predicted and true contact maps, and their insulation profiles (**Additional file 2: Fig. S4B**). Selected flat windows were then evaluated for GC content, ChromHMM states (using the [6] annotation, lifted over to mm10 [58,59]), and their presence in test, training, or validation sets. Following this procedure, we identified 355 flat mouse genomic regions suitable for downstream synthetic boundary insertion experiments. Finally, prediction windows were centered on each flat region such that the midpoint of the flat segment aligned with the midpoint of the contact map predicted by AkitaPT.

### Ledidi Input and Output Loss

Following the original Ledidi [26] framework, the input loss was defined as the total number of nucleotide edits introduced into a sequence. The output loss was calculated in two complementary ways: (1) *global loss*, computed as the L1 distance between the upper-triangular entries of the predicted and target contact maps across the entire window, and (2) *local loss*, computed as the L1 distance restricted to a masked region corresponding to the specific genome-folding feature of interest. The overall optimization objective was defined as a weighted sum of these components:

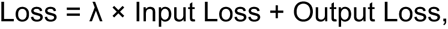

where λ balances the trade-off between minimizing the number of sequence edits and maximizing fidelity to the target contact map. We also tested optimization with the output loss specified as either L2 or Huber loss, but did not observe any gain in design success or performance.

### Map-to-Map Transformations

For proof-of-concept map-to-map transformations, the full AkitaPT model was used with Ledidi. Sequence optimization was performed with a maximum of 2,000 iterations (no early stopping), a default loss weighting of λ=0.01, and edits allowed across the entire input sequence. A global L1 loss was employed to compare predicted and target contact maps, while all other optimization parameters were maintained at their default settings. The similarity between optimized and target maps was quantified using Pearson’s correlation computed on the vectorized upper-triangular entries of the contact maps.

### Akita Semifreddo

#### Memory, Run Time, and Accuracy

We compared four Akita model variants: half-frozen models with 1, 3, or 5 bins of activations replaced after the convolutional tower (the 5-bin model covers the full receptive field) and the full Akita model with all activations recomputed. To simulate a minimal local perturbation, a strong mouse CTCF motif was inserted into a flat genomic region. The relative strengths of natural mouse CTCF motifs were previously characterized in an insertion experiment [33]. This procedure was repeated across 100 naturally occurring strong mouse CTCFs and 24 flat genomic regions, with all metrics averaged over the tested insertions. We reported Pearson’s correlation relative to full Akita outputs, peak memory usage (calculated using torch.cuda.max_memory_allocated(device) on GPU), and prediction speed (**Additional file 1: Table S3**). Additionally, for the full and Akita^SF^ (5-bin) models, memory allocation within each layer was tracked using torch.cuda.memory_allocated() to assess layer-specific resource usage (**Additional file 2: Fig. S3**).

#### Convolutional tower caching

Akita^SF^ is a “half-frozen” version of the AkitaPT model that computes the convolutional tower outputs only for the regions of the input sequence that are being edited (+– receptive field), while using cached activations for the remaining regions. For all 355 flat genomic regions, we first computed and saved the convolutional tower activations as PyTorch .pt tensors for reuse in multiple optimization tasks. Depending on the optimization type:

- For boundary and flame design (edits restricted to central bins), activations for the 5 central bins were recomputed.
- For dot design (edits in two separate bins), activations for the corresponding 10 bins were recomputed.

The recomputed activations were used to overwrite the corresponding slices in the convolutional tower output, and subsequent network layers were then processed as in the full AkitaPT model.

### Genome Folding Features Design

#### Exploring Optimization Parameters

For boundary optimization targeting a mask value of −0.5, we systematically explored a range of Ledidi parameters, including the output loss function (MSE vs. L1), the softmax temperature (τ) used in the straight-through estimator, λ (the trade-off between input and output loss), the learning rate, ε (the finite-difference perturbation size for gradient estimation), the total number of optimization steps, and early stopping criteria. We found that the total number of sequence edits was primarily driven by τ, λ, and ε. Reducing ε substantially decreased the number of edits — from 975 (ε = 1e-3) to 442 (ε = 1e-9) — although further reductions produced diminishing returns. Adjusting τ allowed us to reach a reasonable edit count (∼47.3), but often resulted in faint boundaries. By contrast, λ provided the most interpretable and robust control over edit number: larger λ values consistently reduced the number of edits while maintaining good target fitting. We evaluated λ values between 0.01 and 200, with the final choice yielding ∼50 edits per bin. The same exploration of λ was applied across all feature design tasks.

Based on these observations, we recommend that users begin with the largest λ value that does not substantially increase the optimization failure rate. If fewer edits are desired, λ can be increased further; however, this may necessitate additional optimization steps and a larger number of initial seeds to ensure convergence.

#### Boundary Design

Sequences were optimized using a mask with constant values that directly overwrote the upper right and lower left corners of the target contact map. Optimization parameters were set as follows: max_iter = 2,000 (no early stopping), λ = 125, and edits were restricted to the central bin (index 256). A local L1 loss was applied over the masked region of the map, while all remaining optimization parameters were kept at their default settings.

#### Successful Optimization

An optimization was considered successful if it introduced at least one edit to the input sequence and produced a detectable change in the predicted contact map.

#### Data-driven Dot Mask

A data-driven dot mask was constructed using genomic dots identified by MUSTACHE [60] at 10 kb resolution. Starting from 9,674 candidate dots, we applied the following filters: dots were required to lie on the same chromosome, be located on autosomes, have anchors separated by no more than 384 × 2,048 bp (∼¾ of the predicted map size), and contain CTCF motifs overlapping both anchors in the +– orientation. This filtering yielded 5,713 dots. For each dot, we computed the predicted contact frequency map and extracted a 15 × 15 bin patch centered on the dot, chosen to match the kernel size used by MUSTACHE (∼30kb for 10 kb resolution data). To obtain a representative dot pattern for downstream sequence optimization, we performed a pile-up average using 488 dots with inter-anchor distances of 30-50 bins. The resulting averaged patch was used as the dot mask in all dot-design tasks.

#### Dot Design

Dot design used the data-driven dot mask positioned at the desired inter-anchor distance and integrated into the target map by overwriting the corresponding region. Optimization hyperparameters were tuned for an inter-anchor distance of 50 bins, ensuring that the dot fell well within the ≥100-bin flat genomic regions used for design. Using λ = 130 produced approximately 108.2 edits per optimization, corresponding to ∼54.1 edits per anchor bin. All optimizations were performed with max_iter = 2,000 (no early stopping), λ = 150, and edits restricted to the bins overlapping the target dot anchors. A local L1 loss function was computed over the masked region, and all other optimization parameters were kept at their default values.

#### Flame Design

For flame feature optimization, we applied a stripe-shaped mask (3-bin width, constant value 1.0) and integrated it into the target map by overwriting the corresponding region. All optimizations were performed with max_iter = 2,000 (no early stopping), λ = 140, and edits restricted to the central bin. A local L1 loss function was evaluated over the masked region, and all remaining optimization parameters were left at their default values.

### Feature-specific Scores

#### Boundary Score

Insulation strength was quantified as the mean predicted signal within the upper-right quarter of the contact map. For boundaries designed at the map center, contact depletion is expected in this region; thus, lower boundary scores correspond to stronger boundary formation.

#### Dot Score

Dot strength was measured as the mean predicted signal within a 15 × 15 bin region centered on the intersection of the two dot anchors. Higher dot scores indicate stronger dot features.

#### Flame Score

Flame strength was defined as the mean predicted signal within the 3-bin-wide stripe corresponding to the flame output mask, representing the expected region of contact enrichment.

### Natural Features Strength Analysis

#### Natural Boundaries

We analyzed 4,474 chromatin boundaries previously characterized in the AkitaV2 study [33], identified at 10 kb resolution. For each boundary, predicted contact maps were generated using the mESC [35] fine-tuned AkitaPT model, and the mean boundary score was computed as a quantitative measure of boundary strength.

#### Natural Dots

We analyzed 9,674 chromatin interaction “dots” identified at 10 kb resolution. Dots were excluded if they were located on different chromosomes, on sex chromosomes, or had inter-anchor distances greater than 786 kb (∼three-quarters of the genomic span predicted by Akita). We further restricted the dataset to interactions whose anchors overlapped CTCF motifs in the convergent orientation, yielding a final set of 5,713 dots for downstream analysis. To estimate dot strength, the second anchor was fixed at bin 384 of the Akita prediction map, while the first anchor was positioned according to the original inter-anchor distance. Contact maps were then predicted using the mESC [35] fine-tuned AkitaPT model, and dot strength was quantified using the dot score.

#### Natural Flames

Flame features were detected using StripeNN [42] on the Hsieh mESC Hi-C dataset [35] at 8,192 bp resolution, with default settings. StripeNN initially identified 3,631 flames. To exclude false positives arising from adjacent missing bins, we calculated the fraction of missing bins within a rectangle expanded by 5 bins on all sides of each flame. Bins were considered “bad” if bins.weight was assigned a NaN value during matrix balancing, serving as a proxy for missing data. We retained flames with a missing-bin fraction <0.1 and a positive stripiness score (as recommended by StripeNN), resulting in 2,215 high-confidence flames. To estimate natural flame strength, predicted contact maps were centered at the flame’s lower-left corner (used as the midpoint of the predicted map), and the mean signal within the flame rectangle, as well as the 75th percentile of signal, were computed. For downstream analyses, we selected 1,485 flames with positive mean signal and quantified their strength using the 75th percentile of the signal.

### Sequence Analysis

#### GC Content Analysis

GC content was calculated as the fraction of guanine (G) and cytosine (C) bases within the analyzed sequence. GC content was computed only for the edited portion of the input sequence.

#### CTCF Detection

CTCF motifs were identified using the memesuite-lite library [61,62]. Specifically, the fimo function was employed to scan sequences with the JASPAR [63] CTCF position weight matrix (MA0139.1). All detection parameters were set to their default values.

#### CTCF Pattern Similarity

To compare two sets of CTCF motifs, both genomic positions and orientations were considered. Two motifs were defined as overlapping if they shared the same position and orientation. Pattern similarity was then quantified using the Jaccard index, calculated as the number of overlapping motifs divided by the total number of unique motifs across the two sets.

#### Pre-optimization Motif Score

A motif score for individual 19 bp sites was computed as follows. Positions of 19 bp sites that became CTCF motifs after optimization for strong boundaries were collected from 1,556 successful optimizations, yielding 4,984 sites. Background nucleotide probabilities were estimated from all 355 genomic sequences (each 1.3 Mb in length) containing predicted flat regions at the center. The JASPAR [63] CTCF position weight matrix (MA0139.1) was used to quantify how closely the subsequence of each site matches the expected motif composition relative to the genomic background. For each base, the log₂ ratio between the position weight matrix (PWM)-specified probability of observing that base and its background probability was computed, with a small pseudocount (1 × 10⁻⁹) added to the numerator to avoid undefined logarithms. These values were then summed across all positions of the subsequence to yield the motif score.

#### Aggregated Positive Motif Score

A region-wide aggregated positive motif score was computed for 2,048 bp sequences to provide an estimate of pre-existing CTCF-like information within an edited region. Background nucleotide probabilities were estimated from all 355 genomic sequences (folds 0-7, each 1.3 Mb in length) containing predicted flat regions at their centers. A moving motif similarity score was calculated along the edited portion of each sequence to produce a track of motif-like information content, and positive motif score values were summed across the sequence.

### Transcription Initiation Signal Predictions

Transcription initiation signals were predicted using the Puffin-D model [47]. For each original and edited sequence, predicted GRO-cap signals were obtained for both forward and reverse strands, and the maximum value across strands was used as the maximum predicted GRO-cap signal. Puffin-D predictions were made for 100 kb windows centered on the edited bin or annotated TSS; however, signal quantification was restricted to a smaller region: the editable bin (2,048 bp) for original and designed sequences, and ±1 kb of the annotated TSS coordinate for mouse TSSs. The same procedure was applied to PRO-cap and FANTOM CAGE Puffin-D predictions.

#### Mouse Transcription Start Sites (TSSs)

Mouse TSSs were defined using the GENCODE M25 annotation [64] (available at [65]), considering only entries labeled as “transcript,” where each transcript defines a single isoform with annotated start and end positions. For transcripts on the positive strand, the TSS was assigned to the annotated start coordinate; for negative-strand transcripts, the annotated end was used. To obtain a non-redundant set of TSSs, multiple isoforms per gene were collapsed by selecting the most upstream TSS per strand (smallest coordinate for “+” strand, largest for “– “ strand). After excluding transcripts on sex chromosomes and mitochondrial DNA, this procedure yielded 51,061 unique TSSs from an initial set of 135,182 transcripts.

### Boundary Design Without CTCF Motifs

To prevent the optimization from introducing new CTCF motifs, we incorporated an additional CTCF penalty term into the optimization loss. The total loss function was defined as:

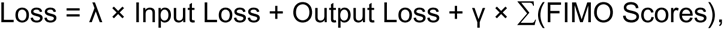

where FIMO scores correspond to all motif occurrences detected using the fimo function from the memesuite-lite library [62], with a default p-value threshold of 10⁻⁴. Under this setup, the optimization objective was to generate a weak boundary (target value = −0.2) while simultaneously suppressing the formation of strong CTCF motifs. The weighting parameter γ was set to 3,000, such that a single strong CTCF motif (FIMO score ≈ 20) would contribute a penalty of ∼60,000 — approximately 2–3× the typical total structural loss (20,000–50,000) — making CTCF introduction strongly unfavorable throughout optimization. The input loss weighting parameter λ was set to 0.01 (default). Optimizations were considered successful if the predicted boundary score of the edited sequence decreased by at least 0.005 relative to the original sequence and no CTCF motifs were detected in the edited bin by FIMO. To assess whether boundary formation was driven by sequence composition rather than specific motifs, optimized sequences were compared against dinucleotide-shuffled controls generated using SeqPro’s k-mer-preserving shuffle [66] with k = 2, which randomizes sequence order while preserving dinucleotide frequencies.

### Suppressing Strong Boundaries

The goal of this analysis was to weaken an existing strong boundary while preventing any edits to CTCF motifs. We reversed the boundary-generation setup: sequences previously optimized to contain a strong (target = −0.5) synthetic boundary at the center of an otherwise flat genomic window were used as inputs, and their corresponding unmodified genomic windows served as targets. CTCF motifs in the synthetic boundary sequences were identified using the fimo function from the memesuite-lite library [62] (default p-value threshold 1 × 10⁻⁴). To preserve these motifs during optimization, Ledidi’s input_mask parameter was used to prevent edits within each CTCF site and ±15 bp flanking regions: for every nucleotide position covered by the mask, all alternative nucleotide weights were strongly suppressed except for the original base, effectively forcing the optimizer to retain the native sequence at those positions. Optimization was performed for 2,000 iterations without early stopping, with λ = 0.01 (default), restricting edits to the central bins and applying a local loss function over the masked region. Boundary suppression was quantified as the change in boundary score before and after optimization, such that positive values indicate successful suppression.

#### Per-bin GC Content

GC content was computed for each 2,048 bp bin across the AkitaPT training set (6,038 sequences of 1,310,720 bp, comprising 640 bins each) and compared to the GC content of the centrally edited bins in boundary-suppressed sequences.

### SINE B2 Analysis in Boundary Suppression

#### SINEB2 Sequence Selection

Three sequence groups were analyzed: isolated strong CTCF motifs, SINE B2 elements overlapping CTCF motifs (SINE B2+CTCF), and SINE B2 elements lacking CTCF motifs (SINE B2 no CTCF). The strong CTCF motif set was defined in our previous work [33] as non-overlapping CTCF motifs non-associated with repetitive elements that overlap boundaries at 10 kb resolution; the 300 motifs with the highest insertion scores were selected. SINE B2 elements were obtained from the UCSC RepeatMasker table for mm10 (https://genome.ucsc.edu/cgi-bin/hgTables?db=mm10&hgta_table%20=%20rmsk, families B2_Mm2, B2_Mm1t, B2_Mm1a; n = 138,585 total). As B2_Mm2 is the most abundant family (n = 94,706), subsequent analyses focused exclusively on this family. B2_Mm2 element coordinates were intersected with JASPAR [63] mm10 CTCF motifs (MA0139.1), yielding 51,779 SINE B2+CTCF and 47,165 SINE B2 no CTCF elements; 300 elements were randomly selected from each group. All analyzed sequences were padded with flanking background sequence to a uniform length of 220 bp, corresponding to the length of the longest elements in the dataset.

#### Background Sequence Generation

To provide a neutral flanking sequence for padding the analyzed elements to a uniform length, we generated background sequences by shuffling 50 selected mouse genomic regions with varying GC content. Shuffling was performed using SeqPro’s k-mer-preserving shuffle [66], which randomizes the sequence while maintaining local k-mer composition. We used k = 8, as this value provided optimal performance in our previous work [33]. Each shuffled candidate sequence (1.3 Mb) was one-hot encoded and passed through the AkitaPT model to predict its contact map. Candidate sequences were evaluated based on two metrics: (1) SCD score, defined as the root-sum-of-squares of the predicted contact map values, capturing overall contact enrichment; and (2) total variation, computed as the sum of absolute differences between adjacent bins along both axes, quantifying local structural variation. Only sequences with SCD and total variation below predefined thresholds (30 and 1300, respectively) were retained, ensuring minimal genome folding signal. For each of the 50 original regions, multiple shuffling attempts were performed using SeqPro until sequences meeting both criteria were obtained. This procedure generated a set of background sequences that preserve base composition and GC content while minimizing predicted folding, providing a neutral baseline for downstream sequence design experiments.

#### First Convolutional Layer Analysis

Each 220 bp sequence was passed through the first convolutional layer of AkitaPT (model.conv_block_1), and the maximum activation of each of the 128 filters was recorded per sequence and per group. Filters that were strongly activated by SINE B2+CTCF elements were identified, and their learned sequence preferences were compared against the B2_Mm2 consensus sequence to assess structural correspondence.

#### Fourth Convolutional Layer Analysis

The same 220 bp sequences were passed through the model up to the fourth convolutional layer (model.conv_tower_block3), whose receptive field of ∼135 bp is sufficient to capture a SINE B2 element. Maximum filter activations were recorded for each sequence. Filters were ranked by SINE B2+CTCF specificity, defined as high activation in the SINE B2+CTCF group relative to both the SINE B2 no CTCF and isolated CTCF groups; filters 68 and 81 exhibited the highest specificity scores. The same procedure was applied to sequences before and after boundary-suppression optimization. Filters 68 and 81 ranked 1st and 5th among filters most strongly upregulated upon optimization, ranked by the difference in mean activation before and after optimization.

#### B-box Shuffling

To evaluate the functional contribution of the Pol III promoter elements, we scanned the boundary-suppression sequences for the B-box consensus sequence (RGTTCRNRTCC), allowing a maximum of three mismatches. The identified matching sequences were then randomly shuffled. These modified sequences were subsequently used as input for AkitaPT to predict changes in boundary strength.

#### SINE B2 Insertion Experiment

To directly test whether SINE B2 sequences are sufficient to suppress CTCF-mediated insulation, one, two, or three copies of the B2_Mm2 consensus sequence were inserted into the central bin of pre-optimization sequences harboring strong synthetic boundaries. Insertion positions were selected randomly within the central bin, subject to the constraint that no insertion overlapped a CTCF motif or its ±15 bp flanking regions. Where multiple copies were inserted, sites were required to be non-overlapping. AkitaPT predictions were generated for each original and modified sequence, and boundary score changes were computed as described above.

#### Relative Suppression Score

The relative suppression score was defined as the ratio of suppression strength to initial boundary strength. Suppression strength was calculated as the change in boundary score after versus before suppression. Boundary strength was defined as the difference between the boundary score at the initial boundary and that of a flat region prior to optimization. A value of 0 corresponds to the initial boundary, and 1 indicates complete suppression to the flat-region level.

### In-silico Validation

To validate our designed sequences, we used AlphaGenome [30] as an independent *in silico* evaluation tool. AlphaGenome was chosen because it (1) was developed independently of Akita and (2) predicts genome folding at the same 2,048 bp resolution.

If not stated otherwise, we specified the ontology term EFO:0004038, corresponding to *in situ* Hi-C data from mouse embryonic stem cells (mESCs) [67], matching the cell type used to fine-tune AkitaPT. For each designed sequence, the bins corresponding to the modified regions were replaced with the optimized sequence, and the resulting change in feature strength was quantified. Differences in predicted feature strength between the original and optimized sequences were then calculated using AlphaGenome, and concordance with AkitaPT predictions was assessed using Pearson’s correlation coefficients.

#### Contact map RMSD

For dot designs, the localized dot score showed moderate cross-model agreement. To provide a more global measure of map change, we additionally computed the contact map RMSD between original and optimized sequence predictions. RMSD was defined as the root mean squared pixelwise difference over all upper-triangular entries of the contact map, excluding the two nearest off-diagonals.

### Designs displaying limitations of the approach

#### B-cell Model Training

Fountains are not detectable in all cell types; therefore, we trained a model on a cell type in which fountains are visibly apparent. We collected Hi-C data from B cells via the 4DN data portal [43], specifically the paired files 4DNFI27I3P1V and 4DNFIFBBAKK4, which were concatenated to increase coverage. Data processing followed the protocol described in [33], with modifications for enhanced smoothing: the standard deviation of the Gaussian2D kernel was increased to 2, and the adaptive coarse-graining cutoff was set to 4.

Eight models were trained from scratch on distinct data splits using the schedulefree AdamW optimizer [54] with the following hyperparameters: BATCH_SIZE = 4, EPOCHS = 200, LR = 0.01, L2_SCALE = 1.5 × 10⁻⁵, WEIGHT_CLIPPING = 20.0, and EARLY_STOP = 50. Training progress was monitored to ensure loss convergence. Despite the limited quality of the data, the models achieved relatively good predictive performance, as assessed by Pearson’s r = 0.58, Spearman’s r = 0.51, and MSE = 0.12 (average over an ensemble of 8 models).

#### Fountain Design

For fountain feature design, we defined a fountain mask as a symmetric, antidiagonal cone-shaped region, widest at the top-right and bottom-left corners (up to 120 bins), tapering linearly to a single point at the antidiagonal midpoint. A constant mask value of 0.5 was applied and integrated into the target map via overwriting. We employed a multi-model optimization setup, providing four models with their corresponding target maps, while using the remaining four models for validation. The output loss was computed as the sum of local L1 losses over the masked region between each model’s predicted map and its respective target. Optimization was performed for 2,000 iterations without early stopping, with default λ = 0.01, and edits restricted to the central 50 bins.

#### Cell-Type Specific Boundary Design

For cell-type-specific boundary designs, we utilized two human micro-C datasets (H1hESC and HFF) [68], as AlphaGenome [30] supports predictions across multiple human cell types. We employed a multi-model approach using four Akita^SF^ models: two fine-tuned on H1hESC data and two fine-tuned on HFF data. The Akita^SF^+Ledidi framework was provided with the initial sequence and four corresponding target contact maps, one for each model. The optimization loss was calculated as the sum of local L1 losses across the masked region, comparing each model’s predicted contact map to its respective target. Optimization proceeded for 2,000 iterations without early stopping, using a regularization parameter of λ = 0.01 (default). For *in silico* validation of cell-type-specific designs using AlphaGenome, we employed ontology terms EFO:0003042 and EFO:0009318, corresponding to H1hESC and HFF cell types, respectively.

### Statistics and Software

The neural network was implemented in Python (v3.10) using the PyTorch (v2.3) framework. Statistical significance tests are specified in the main text and relevant figure legends.

Correlation coefficients (Pearson’s r and Spearman’s r) were calculated using SciPy (v1.15) [69]. Data processing and visualization were performed using NumPy (v1.26.4) [70], pandas (v2.3.3) [71], Matplotlib (v3.10.8) [72], and Seaborn (v0.13.2) [73].

## Code Availability

All code required to reproduce the results is publicly available. Scripts for Akita-PyTorch fine-tuning and training can be found at https://github.com/PSmaruj/pytorch_akita; model weights are deposited at https://doi.org/10.5281/zenodo.19599537. The code for Akita^SF^ + Ledidi optimization and subsequent analysis is hosted at https://github.com/PSmaruj/akita_semifreddo.

## Acknowledgements

G.F. and P.N.S. are supported by the National Institute of General Medical Sciences R35 GM143116-01. The authors would like to thank Jacob Schrieber for helpful discussions about Ledidi, the Fudenberg group for general feedback, and the contributing artists at OpenClipart (https://openclipart.org/) for providing the water puddle and ice cube vector icons under a free-use license.

## Notes

https://github.com/Fudenberg-Research-Group/akita_semifreddo

https://github.com/Fudenberg-Research-Group/akita_pytorch

### Competing Interest Statement

D.R.K. is an employee of Calico Life Sciences, LLC. All other authors declare no competing interests.

