## Supplemental Information for "Sequence design for three-dimensional genome folding using Akita Semifreddo"

#### Additional file 1: Supplementary Tables S1–S9

| Cell type | Model | Pearson's r | Spearman's r | MSE |
|---|---|---|---|---|
| mESC | Tensorflow original | 0.638 | 0.583 | 0.124 |
| | PyTorch transferred | 0.588 | 0.529 | 0.140 |
| | PyTorch fine-tuned | 0.663 | 0.613 | 0.117 |
| HFF | Tensorflow original | 0.667 | 0.606 | 0.176 |
| | PyTorch transferred | 0.627 | 0.556 | 0.223 |
| | PyTorch fine-tuned | 0.674 | 0.613 | 0.177 |

**Additional file 1: Table S1. Comparison of Akita models.**
This table compares three versions of the Akita model: the original AkitaV2 TensorFlow model [33], a PyTorch implementation with transferred TensorFlow weights, and the PyTorch model after framework-specific fine-tuning. Results are shown for one mouse model (mESC, [35]) and one human model (HFF, [68]). Values represent averages over eight models trained on different data splits. Pearson's r, Spearman's r, and MSE were calculated by flattening all target and predicted matrices into concatenated vectors and computing metrics between them. Fine-tuned PyTorch models exhibit slightly improved performance relative to the original TensorFlow models, likely reflecting the use of separate models for each cell type, whereas AkitaV2 was trained to predict multiple cell types simultaneously.

| Species | Cell type | $Akita_{PT}$ | AlphaGenome [30] | ORCA [34] |
| --- | --- | --- | --- | --- |
| mouse | mESC [35] | r: 0.654<br>ρ: 0.602<br>MSE: 0.119 | | |
| | mESC [67] | r: 0.748<br>ρ: 0.700<br>MSE: 0.077 | r: 0.667<br>ρ: 0.621<br>MSE: 0.101 | |
| | cortical neurons [67] | r: 0.797<br>ρ: 0.747<br>MSE: 0.076 | | |
| | neocortex cortical neuron [67] | r: 0.770<br>ρ: 0.716<br>MSE: 0.078 | | |
| | neural progenitor cell [67] | r: 0.780<br>ρ: 0.728<br>MSE: 0.071 | | |
| | neocortex neural progenitor cell [67] | r: 0.735<br>ρ: 0.676<br>MSE: 0.068 | | |
| human | HFF [68] | r: 0.675<br>ρ: 0.615<br>MSE: 0.176 | r: 0.724<br>ρ: 0.672<br>MSE: 0.162 | r: 0.508<br>ρ: 0.465<br>MSE: 0.295 |
| | H1hESC [68] | r: 0.683<br>ρ: 0.642<br>MSE: 0.138 | r: 0.726<br>ρ: 0.691<br>MSE: 0.120 | r: 0.605<br>ρ: 0.584<br>MSE: 0.182 |
| | GM12878 [41] | r: 0.631<br>ρ: 0.589<br>MSE: 0.102 | r: 0.652<br>ρ: 0.603<br>MSE: 0.103 | |
| | IMR90 [41] | r: 0.637<br>ρ: 0.579<br>MSE: 0.197 | r: 0.672<br>ρ: 0.616<br>MSE: 0.192 | |
| | HCT116 [74] | r: 0.652<br>ρ: 0.601<br>MSE: 0.081 | r: 0.598<br>ρ: 0.556<br>MSE: 0.099 | |

**Additional file 1: Table S2. Benchmarking of released models.**
Performance for six mouse cell types ($Akita_{PT}$ models and AlphaGenome) and five human cell types ($Akita_{PT}$, AlphaGenome, and ORCA). Cells corresponding to datasets on which a model

was not trained, and corresponding predictions are not available, are shaded black. For mouse predictions, overlapping test datasets between $Akita_{PT}$ and AlphaGenome yielded 2,940 windows, from which 500 were randomly selected for performance evaluation. For human predictions, overlapping test datasets among $Akita_{PT}$, AlphaGenome, and ORCA yielded 352 windows. Pearson's r, Spearman's r, and MSE were calculated by flattening all target and predicted matrices into concatenated vectors. Fine-tuned Akita PyTorch models achieve comparable performance to AlphaGenome and ORCA, except on Micro-C data, where AlphaGenome performs better, likely reflecting its multi-modal training on additional data such as accessibility and TF binding.

| Number of bins replaced | Pearson’s r | Memory peak [MB] | Time [ms] |
|---|---|---|---|
| 1 | 0.9886 | 582.804 | 3.938 |
| 3 | 0.9997 | 582.804 | 3.903 |
| 5 ($Akita^{SF}$) | 1.0 | 582.804 | 3.944 |
| All ($Akita_{PT}$) | 1.0 | 1735.965 | 5.052 |

**Additional file 1: Table S3. Comparison of memory usage and prediction speed between Akita Semifreddo and full Akita model.**

Comparison between the full $Akita_{PT}$ model and semi-frozen variants. In all semi-frozen cases, 11 bins were recomputed to span the entire receptive field around the edited central bin and eliminate padding artifacts. We evaluated versions where 1, 3, or 5 bins were replaced following the convolutional tower (the 5-bin version representing the full receptive field). To simulate a minimal local change, a strong mouse CTCF motif was inserted into the central bin of a flat genomic region (**Methods**). This procedure was repeated across 100 naturally occurring strong mouse CTCFs and 24 flat genomic regions, each averaged over all tested insertions. Columns show Pearson’s r (compared to full Akita outputs), average peak memory usage, and prediction time. Recomputing 5 bins yields outputs identical to the full model, while reducing peak memory usage threefold and improving speed by ~25%.

| λ | No-edit optimizations | Edits present, no contact-depletion optimizations | Mean edits (successful only) | Mean boundary score difference (successful only) | Mean CTCF motif count (successful only) |
|---|---|---|---|---|---|
| 0.01 (default) | 0 / 174 | 0 / 174 | 792.098 | -0.314 | 7.121 |
| 0.1 | 0 / 174 | 0 / 174 | 757.385 | -0.315 | 7.063 |
| 1.0 | 0 / 174 | 0 / 174 | 430.977 | -0.306 | 6.655 |
| 10.0 | 1 / 174 | 0 / 174 | 226.253 | -0.302 | 6.092 |
| 100.0 | 14 / 174 | 0 / 174 | 63.006 | -0.218 | 3.270 |
| 125.0 | 18 / 174 | 0 / 174 | 51.115 | -0.202 | 2.839 |
| 200.0 | 29 / 174 | 0 / 174 | 31.943 | -0.169 | 2.144 |

**Additional file 1: Table S4. Selection of the λ parameter for boundary optimization.**

For strong boundary optimization (target = -0.5), λ (the scaling factor between input and output loss) was varied between 0.01 (default) and 200 to achieve approximately 50 edits per editable DNA bin (2,048 bp). λ = 125, yielding ~51 edits, was selected for subsequent optimizations.

| λ | No-edit optimizations | Edits present, no contact-enrichment optimizations | Mean edits (successful only) | Mean dot score difference (successful only) | Mean CTCF motif count (successful only) |
|---|---|---|---|---|---|
| 0.01 (default) | 0 / 174 | 0 / 174 | 1136.333 | 0.557 | 1.856 |
| 0.1 | 0 / 174 | 0 / 174 | 1135.943 | 0.553 | 1.661 |
| 1.0 | 0 / 174 | 0 / 174 | 1036.741 | 0.551 | 1.960 |
| 10.0 | 0 / 174 | 0 / 174 | 596.621 | 0.551 | 1.914 |
| 100.0 | 5 / 174 | 0 / 174 | 169.425 | 0.530 | 2.259 |
| 150.0 | 26 / 174 | 0 / 174 | 108.178 | 0.461 | 2.063 |
| 200.0 | 44 / 174 | 0 / 174 | 48.902 | 0.410 | 2.011 |

**Additional file 1: Table S5. Selection of the λ parameter for dot optimization.**
For dot optimization with an inter-anchor distance of 50 bins, λ was tested between 0.01 and 200. Because edits are allowed across two bins overlapping the dot anchors, λ = 150 was selected, yielding ~108 edits across 2 bins (~54 edits per bin).

| λ | No-edit optimizations | Edits present, no contact-enrichment optimizations | Mean edits (successful only) | Mean flame score difference (successful only) | Mean CTCF motif count (successful only) |
|---|---|---|---|---|---|
| 0.01 (default) | 0 / 174 | 0 / 174 | 744.730 | 0.715 | 4.310 |
| 0.1 | 0 / 174 | 0 / 174 | 737.747 | 0.725 | 4.678 |
| 1.0 | 0 / 174 | 0 / 174 | 672.063 | 0.709 | 4.425 |
| 10.0 | 0 / 174 | 0 / 174 | 376.879 | 0.704 | 4.460 |
| 100.0 | 7 / 174 | 0 / 174 | 80.115 | 0.662 | 3.276 |
| 140.0 | 11 / 174 | 0 / 174 | 58.540 | 0.646 | 2.937 |
| 200.0 | 18 / 174 | 0 / 174 | 41.810 | 0.614 | 2.626 |

**Additional file 1: Table S6. Selection of the λ parameter for flame optimization.**
For flame optimization, λ was tested between 0.01 and 200. λ = 140, yielding ~58.5 edits per bin, was selected for subsequent optimizations.

| ε | No-edit optimizations | Edits present, no contact-depletion optimizations | Mean edits (successful only) | Mean boundary score difference (successful only) | Mean CTCF motif count (successful only) |
|---|---|---|---|---|---|
| 1e-9 | 0 / 174 | 0 / 174 | 441.879 | -0.279 | 5.891 |
| 1e-8 | 0 / 174 | 0 / 174 | 468.052 | -0.285 | 5.925 |
| 1e-7 | 0 / 174 | 0 / 174 | 491.443 | -0.290 | 6.293 |
| 1e-6 | 0 / 174 | 0 / 174 | 530.023 | -0.302 | 6.511 |
| 1e-5 | 0 / 174 | 0 / 174 | 618.431 | -0.308 | 6.563 |
| 1e-4 (default) | 0 / 174 | 0 / 174 | 792.098 | -0.314 | 7.121 |
| 1e-3 | 0 / 174 | 0 / 174 | 975.063 | -0.317 | 7.276 |

**Additional file 1: Table S7. Effect of ε on the number of edits in strong boundary optimization.**

To reduce the number of sequence edits, we varied ε (the perturbation added to logits/weights) from $1\times10^{-3}$ to $1\times10^{-9}$. Across this range, the average number of edits decreased from 975 to 442. However, the trend plateaued at lower ε values, indicating that substantially reducing the number of edits cannot be achieved by tuning ε alone.

| τ | No-edit optimizations | Edits present, no contact-depletion optimizations | Mean edits (successful only) | Mean boundary score difference (successful only) | Mean CTCF motif count (successful only) |
|---|---|---|---|---|---|
| 0.01 | 0 / 174 | 6 / 174 | 47.270 | -0.007 | 0.690 |
| 0.1 | 0 / 174 | 0 / 174 | 360.195 | -0.218 | 4.040 |
| 1.0 (default) | 0 / 174 | 0 / 174 | 792.098 | -0.314 | 7.121 |
| 10.0 | 0 / 174 | 0 / 174 | 1079.172 | -0.319 | 7.626 |

**Additional file 1: Table S8. Effect of τ (Gumbel-Softmax temperature) on the number of edits during strong boundary optimization.**

To test whether the number of sequence edits could be reduced, τ was varied across 0.01, 0.1, 1.0, and 10.0. Although τ = 0.01 yielded a low number of edits (~47 per bin), the resulting changes in boundary insulation were minimal. Thus, tuning τ did not provide a viable strategy for reducing edits while maintaining optimization efficacy.

| Design | Target feature score / distance (in bins) | No-edit optimizations | Success rate [%] | Mean edits (successful only) |
|---|---|---|---|---|
| Boundaries | -0.2 | 38 / 355 | 89.3 | 23.2 |
| | -0.3 | 28 / 355 | 92.1 | 31.4 |
| | -0.4 | 41 / 355 | 88.5 | 42.8 |
| | -0.5 | 32 / 355 | 91.0 | 55.6 |
| | -1.0 | 48 / 355 | 86.5 | 112.8 |
| | -5.0 | 50 / 355 | 85.9 | 124.8 |
| Dots | 30 | 115 / 355 | 67.6 | 222.5 |
| | 50 | 48 / 355 | 86.5 | 127.3 |
| | 70 | 13 / 355 | 96.3 | 111.0 |
| Flames | 0.4 | 44 / 355 | 87.6 | 38.1 |
| | 0.6 | 22 / 355 | 93.8 | 44.3 |
| | 0.8 | 26 / 355 | 92.7 | 55.7 |
| | 1.0 | 23 / 355 | 93.5 | 63.3 |

**Additional file 1: Table S9. Success rates and edit statistics across genome-folding optimization tasks.**

For each genome-folding feature, we performed optimizations across multiple target strengths (boundaries, flames) or inter-anchor distances (dots). For each configuration, we report the number of unsuccessful runs — those in which no edits were accepted or the desired map modification was not achieved — and compute the corresponding success rate. Among successful optimizations, we further report the average number of edits.

### Additional file 2: Supplementary Figures S1–S11

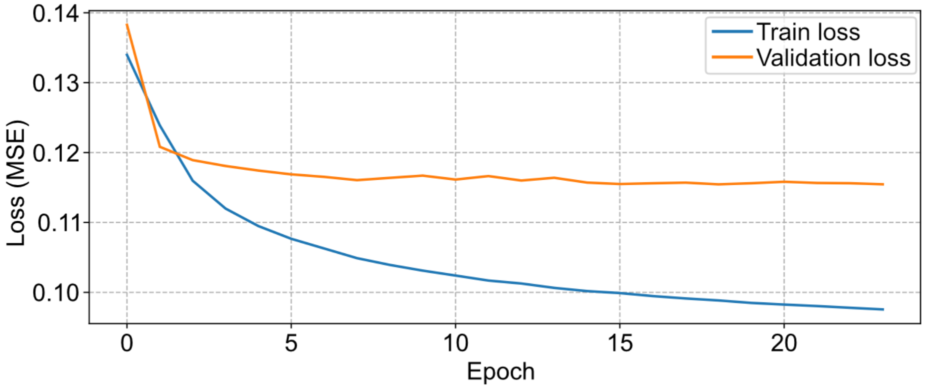


**Additional file 2: Figure S1. Example fine-tuning training curves.**
Shown are training dynamics from framework-transfer fine-tuning (TensorFlow to PyTorch) for one Akita model fine-tuned to mESC data [35]. Training loss (blue) and validation loss (orange) both exhibit a large improvement during the first epoch, followed by rapid stabilization and plateauing over subsequent epochs.

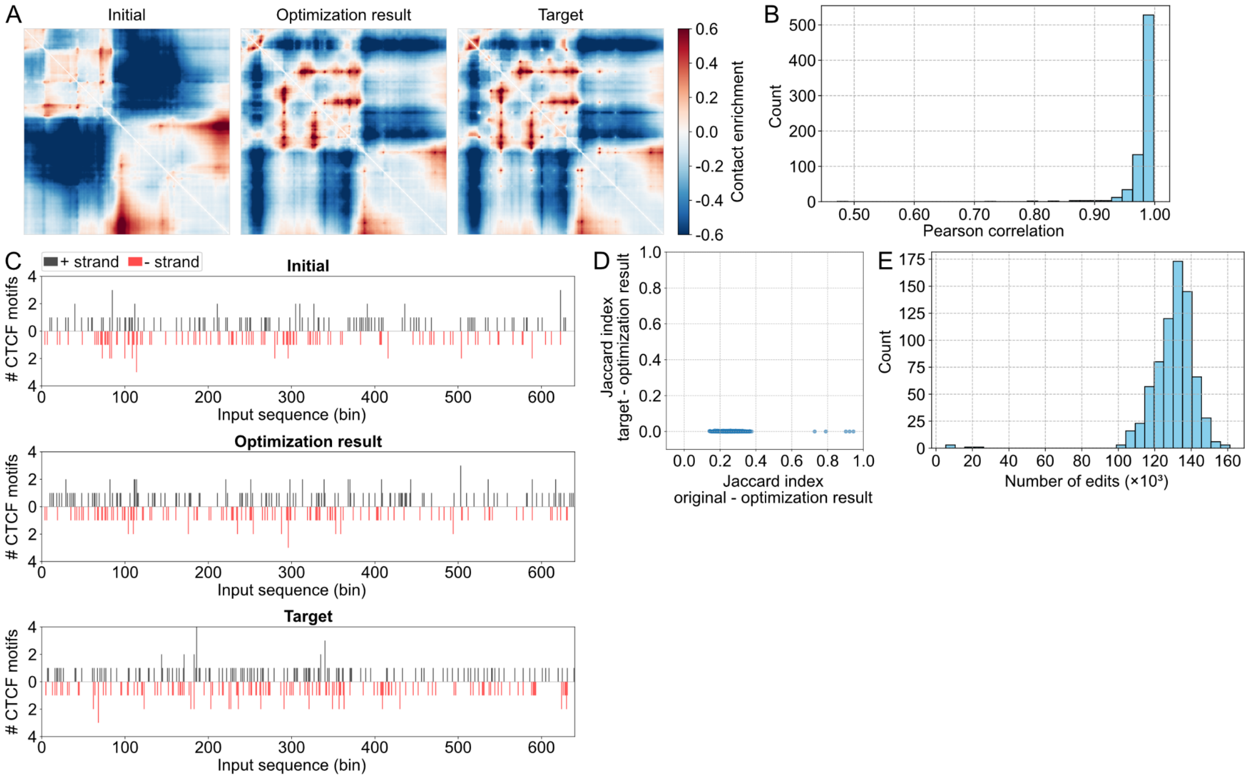


**Additional file 2: Figure S2. Akita$^{PT}$ combined with the Ledidi optimizer can “rewire” genome folding.**

**A)** Three contact maps are shown: the original genome folding prediction, the optimized genome folding prediction, and the target genome folding prediction.

**B)** Histogram of Pearson's correlation between the optimized and target genome folding predictions for 725 genome folding “rewiring” optimizations.

**C)** CTCF motif landscapes for the initial (top), optimized (middle), and the sequence corresponding to the target maps (bottom). The x-axis spans 640 bins (~1.3 Mb). Positive black bars indicate forward-strand CTCF motifs, and negative red bars indicate reverse-strand motifs.

**D)** Overlap fraction of CTCF motifs between the initial and optimized sequences, and between the optimized and target sequences. Motifs were considered overlapping only if they matched in both genomic location and strand. The optimized sequences remain more similar to the original sequences than to those underlying the target folding patterns.

**E)** Histogram showing the number of accepted sequence edits required to achieve map-to-map transformations for 725 genome folding “rewiring” optimizations. Edits were allowed across the entire input sequence (~1.3 Mb).

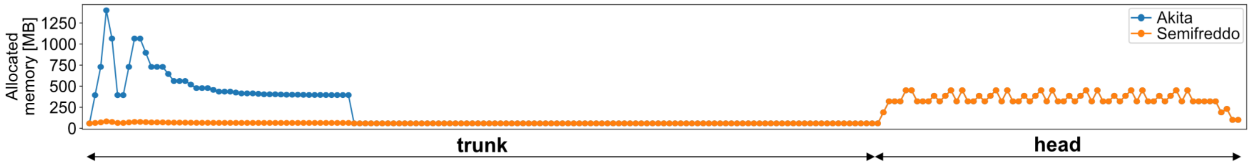


**Additional file 2: Figure S3. Akita$^{SF}$ vs. full Akita memory usage.**
Allocated GPU memory per layer for the full Akita model (blue) and the Akita$^{SF}$ model (orange), averaged over 2,400 predictions with central CTCF insertions. The x-axis indicates layer indices in forward order. Akita$^{SF}$ bypasses the memory-intensive convolutional tower, avoiding the large memory peak.

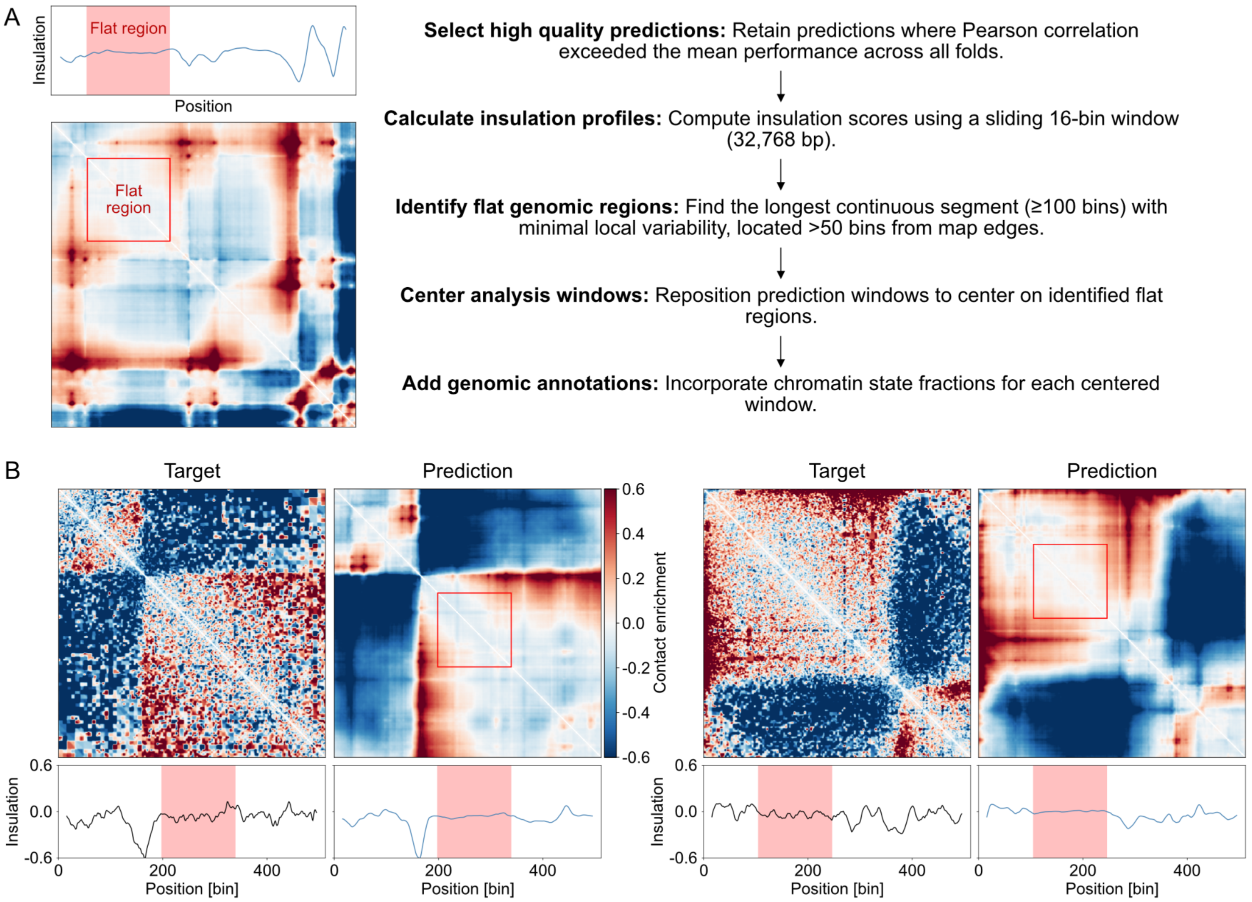


**Additional file 2: Figure S4. Selection of optimization input sequences.**

**A)** Selection of optimization input sequences. We began with a single cross-validation fold (1/8 of the training data). For each prediction window in this fold, we computed Pearson's correlation between the model prediction and the target. Only windows with Pearson's r above the fold-specific average were retained, ensuring that optimization starts from regions the model predicts well. Next, we identified long continuous segments (≥100 bins, ~200 kb) in which the insulation profile showed minimal local variability and which were located away from map edges (>50 bins from either edge). Prediction windows were then centered on these flat regions. For each selected window, chromatin states were computed as in [6].

**B)** Two example target-prediction pairs with detected flat regions highlighted as red rectangles on the predicted maps. Insulation profiles aligned below each map also show the corresponding flat regions (red), indicating where the selection criteria were met.

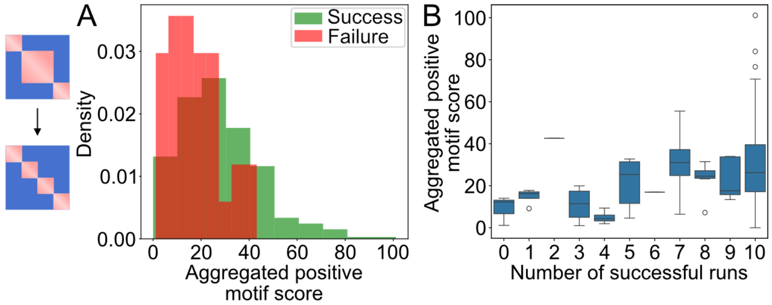


**Additional file 2: Figure S5. Boundary optimization success depends on aggregated positive CTCF motif scores.**

**A)** To understand why strong boundary optimizations failed for some sequences, we computed an aggregated positive motif score, defined as the sum of all positive CTCF motif scores across the entire edited sequence prior to optimization. Comparison of successfully optimized sequences (green) with failed sequences (red) revealed that failed optimizations tend to exhibit lower aggregated positive motif scores. This metric was more predictive of optimization success than other pre-existing sequence features, including: (i) the number of positive motif instances, (ii) the sum of all motif scores (positive and negative), and (iii) the maximum motif score.

**B)** Each of the 174 sequences was subjected to strong boundary optimization across 10 independent runs. Boxplots show the distribution of aggregated positive motif scores as a function of the number of successful optimizations (out of 10 independent runs). Sequences with more frequent optimization success tend to have higher aggregated positive motif scores. Note that only one sequence displayed either 2 or 6 sucessful runs.

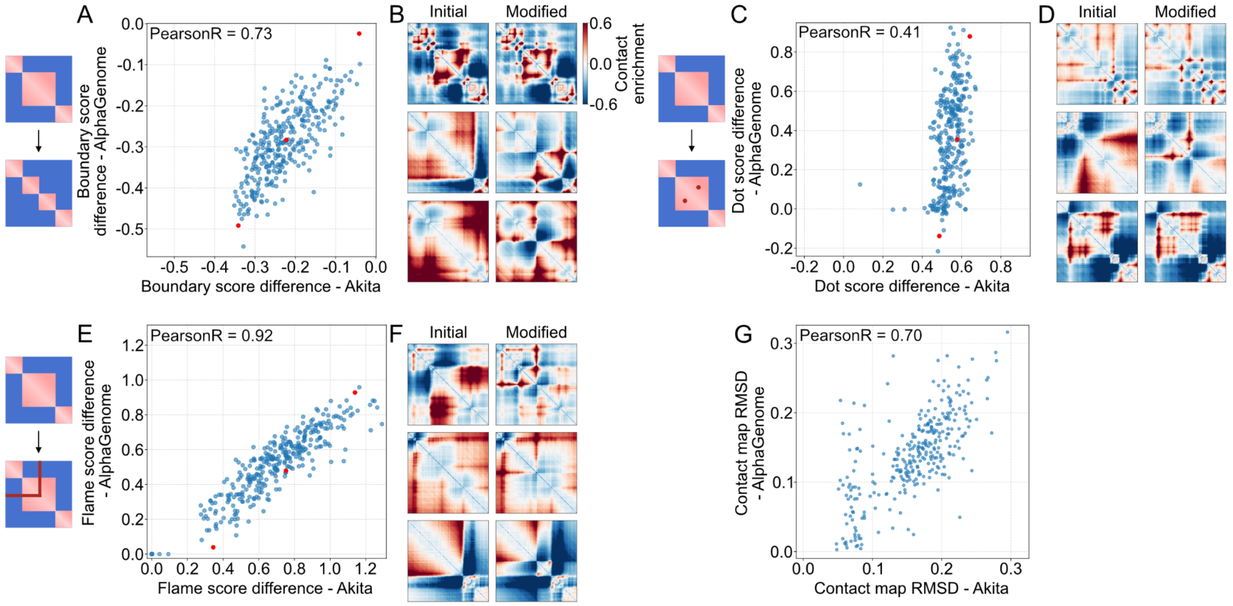


**Additional file 2: Figure S6. Independent validation of boundary, dot, and flame designs with AlphaGenome.**

**A)** Quantitative validation with AlphaGenome for boundary design (target = -0.5). Boundary score differences (optimized – initial) were computed for both Akita and AlphaGenome predictions and show strong agreement (Pearson's r > 0.7).

**B)** Qualitative validation with AlphaGenome. Three examples comparing AlphaGenome predictions for original sequences and sequences with edited central bins that induce boundary formation, providing *in silico* validation of the designed boundaries. The three pairs of plots correspond to the three red points in panel F, from top to bottom.

**C)** Quantitative validation of dot designs at 50 bin distance. Dot score differences (optimized − initial) were computed for both Akita and AlphaGenome predictions. The results show moderate agreement (Pearson's r > 0.4), likely due to the very local nature of this score (see G).

**D)** Three examples of AlphaGenome predictions showing original sequences and the corresponding sequences with dot-inducing edits. Examples are ordered by dot score from highest to lowest (top to bottom). The color scale is the same as in panel B.

**E)** Quantitative validation of flame designs (target = 1.0). Flame score differences (optimized – initial) computed for Akita and AlphaGenome predictions show strong concordance (Pearson's r > 0.9).

**F)** Qualitative validation of flame designs. Three examples of AlphaGenome predictions for original sequences and the same sequences after flame-inducing edits. The color scale is the same as in panel B.

**G)** To further validate dot design quality, contact map RMSD (root mean squared pixelwise difference between original and designed sequence predictions) was computed as a more global measure of map change. While the localized dot score (averaged over a 15×15 pixel patch centered on the dot position) showed moderate cross-model agreement, the contact map RMSD showed substantially higher correlation between Akita and AlphaGenome predictions, indicating that the broader structural changes induced by the designed sequences are consistent across both models.

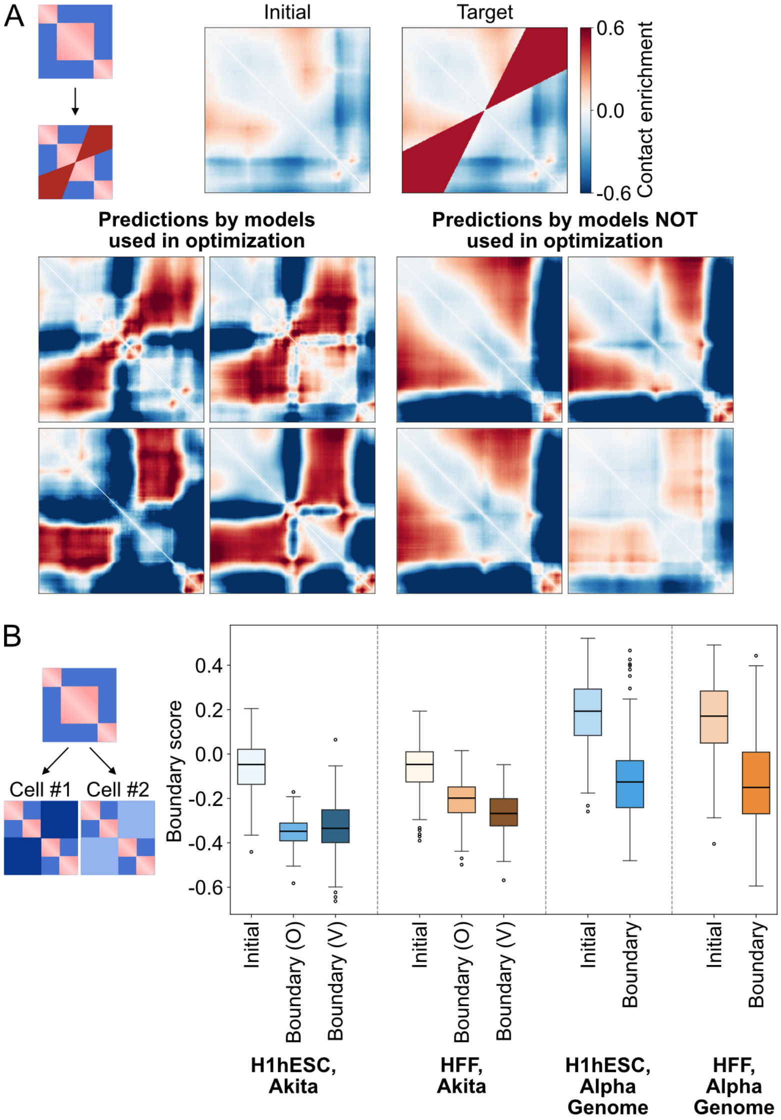


**Additional file 2: Figure S7. Failed validation of fountain designs and cell-type-specific boundaries.**

**A)** Unsuccessful fountain designs using models trained on B-cell data [43], where fountain structures were characterized without data aggregation. *Top:* contact map before optimization showing a flat genomic region in the center, and the target map with a fountain mask applied. *Bottom:* predictions from four models used in the multi-model fountain optimization show visible fountain-like structures, while predictions from four models not used in optimization (serving as validation models) do not exhibit fountain-like structures, precluding visual validation of the designs.

**B)** Unsuccessful cell-type-specific boundary design. The optimization objective was to create a sequence resulting in a strong boundary in H1hESC (blue) and a weak boundary in HFF (orange). Boxplots show the distributions of boundary scores: (i) H1hESC boundary scores before optimization, after optimization as predicted by two Akita models used in optimization, and after optimization as predicted by two Akita models not used in optimization (validation models); (ii) HFF boundary scores before optimization, after optimization as predicted by two Akita models used in optimization, and after optimization as predicted by two validation models; (iii) H1hESC boundary scores before and after optimization as predicted by AlphaGenome; (iv) HFF boundary scores before and after optimization as predicted by AlphaGenome. Both the Akita optimization and validation models show that the designed boundaries are stronger in H1hESC than in HFF. However, the distributions of boundary scores calculated from AlphaGenome predictions overlap substantially, failing to validate the cell-type-specific design.

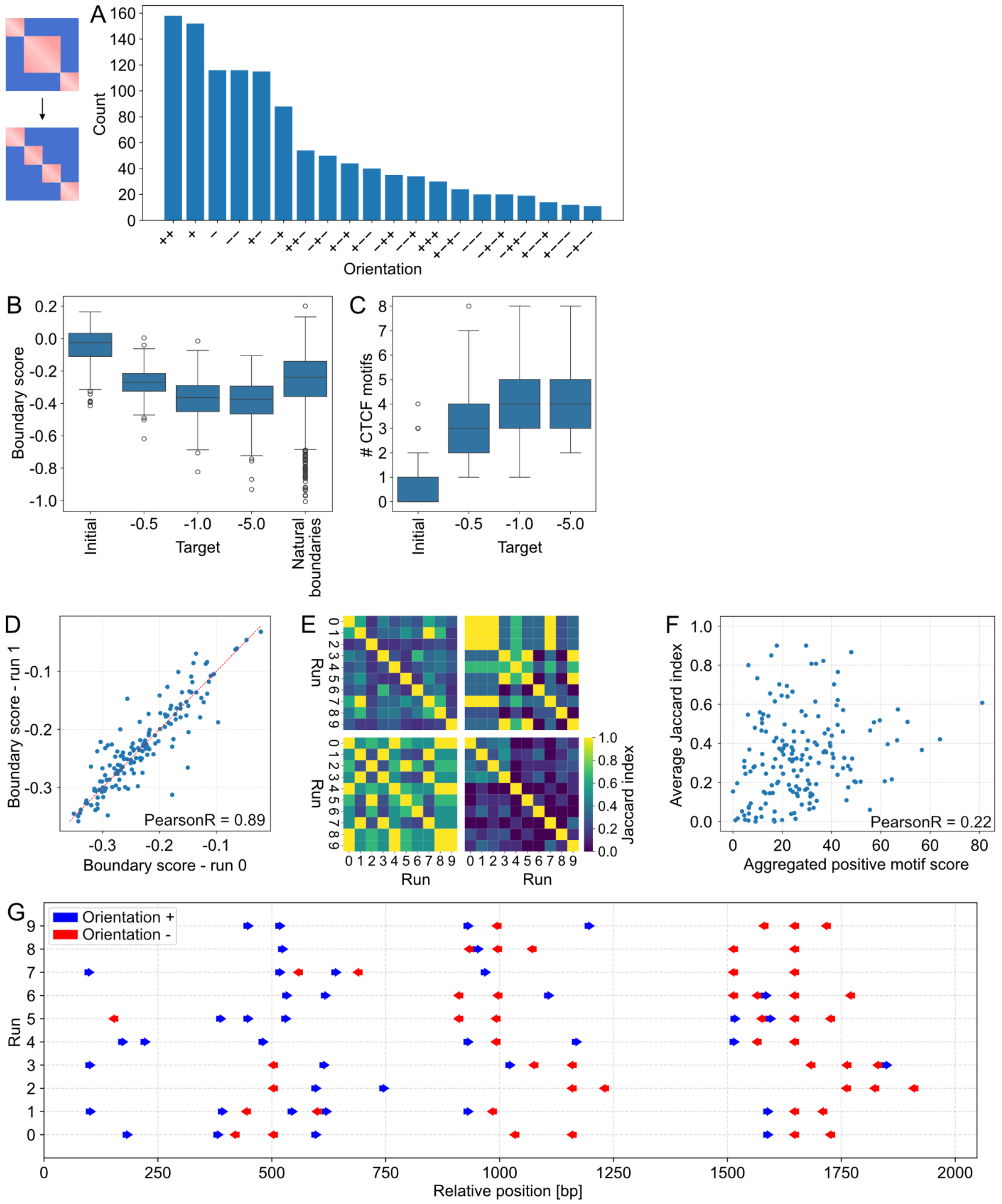


**Additional file 2: Figure S8. Sequence features of designed boundaries.**

**A)** Histogram of CTCF motifs orientations across all boundary optimization experiments (targets -0.2, -0.3, -0.4, -0.5). Only orientations with counts exceeding 10 are shown for clarity.

**B)** Boundary scores for natural boundaries (estimated from >4,000 mouse boundaries), initial genome-folding predictions, and predictions for sequences optimized toward strong (-0.5) and extreme (-1, -5) boundary strengths. Boundary scores after optimization plateau at extreme target values, indicating that such strong boundaries are not achievable within our framework.

**C)** Number of detected CTCF motifs for initial sequences and sequences optimized toward strong (-0.5) and extreme (-1, -5) boundary strengths. The number of CTCF motifs does not scale proportionally with increasingly strong target boundaries.

**D)** Reproducibility of boundary designs. Boundary score differences were calculated for two independent runs of the same optimization (same targets, same genomic regions). The results are highly consistent (Pearson's r > 0.8).

**E)** Four examples of regions for which all 10 independent optimization runs were successful. For each pair of runs, we calculated the Jaccard index between the sets of detected CTCF motifs. Some regions show high consistency across runs, while others yield more diverse solutions.
**F)** For each optimized sequence, we computed the average Jaccard index of CTCF motif sets across all runs versus the aggregated positive motif score prior to optimization. The scatterplot shows that sequences with higher aggregated positive motif scores exhibit higher average Jaccard indices (Pearson’s $r = 0.22$), indicating lower diversity of optimized CTCF configurations. This suggests that optimization tends to reuse pre-existing motif-like features when such features are more abundant in the initial sequence.
**G)** CTCF motif positions detected in ten independent optimizations as in Figure 3D for a strong (target = -0.5) boundary optimizations but performed with a much lower $\lambda = 0.01$. These runs produce more CTCF motifs and exhibit greater diversity in CTCF motifs across independent optimizations.

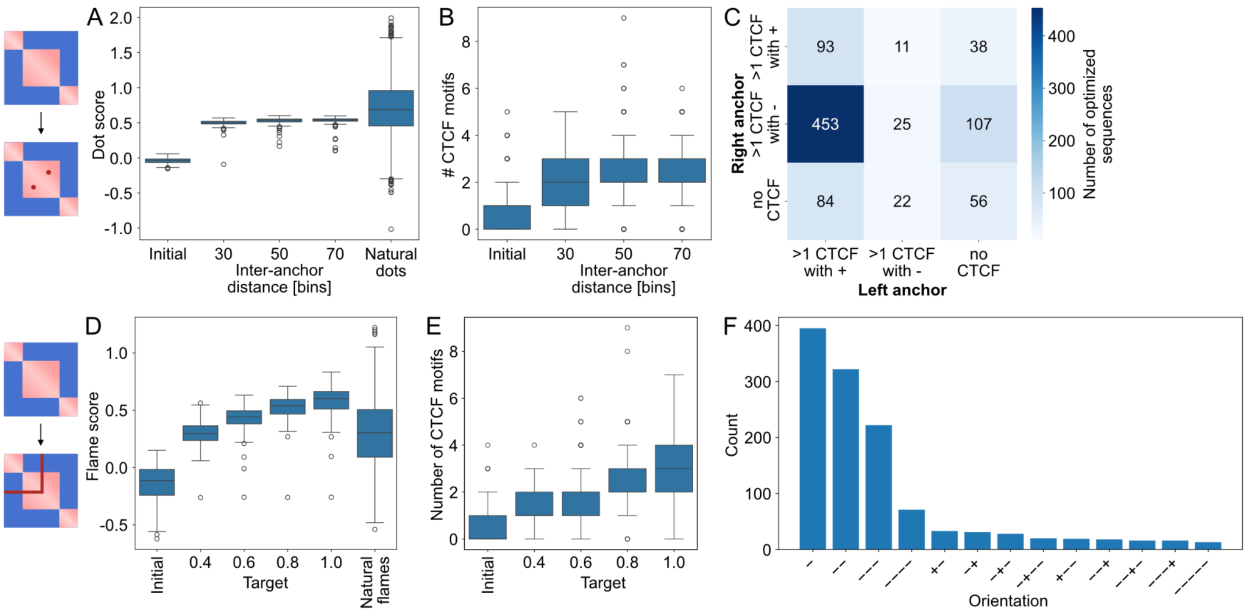


**Additional file 2: Figure S9. Sequence features of designed dots and flames.**

**A)** Dot scores, computed as the average contact frequency within a 15×15 dot mask, are shown for natural dots (estimated from >6,800 mouse dots) and designed dots with inter-anchor distances of 30, 50, 70, and 90 bins.

**B)** Number of detected CTCF motifs in the two editable bins (overlapping the dot anchors) for initial sequences and designed sequences across the four tested inter-anchor distances.

**C)** Classification of CTCF configurations at dot anchors. For each anchor, CTCFs were categorized as: more than one CTCF in the forward orientation, more than one in the reverse orientation, or none detected. The matrix shows combinations of these categories for the left (columns) and right (rows) anchors. The most frequent configuration is +−, consistent with naturally occurring dots [41].

**D)** Flame scores, computed as the average contact frequency within a 3-bin–wide flame mask, are shown for >1,400 natural flames and for four target flame strengths. Designed flame strengths scale with the specified target values.

**E)** Number of detected CTCF motifs within the edited subsequences for the four target flame strengths.

**F)** Histogram of CTCF motifs orientations aggregated across all flame designs. The most frequent orientations are unidirectional (−, −−, −−−). Only orientations with counts exceeding 10 are shown for clarity.

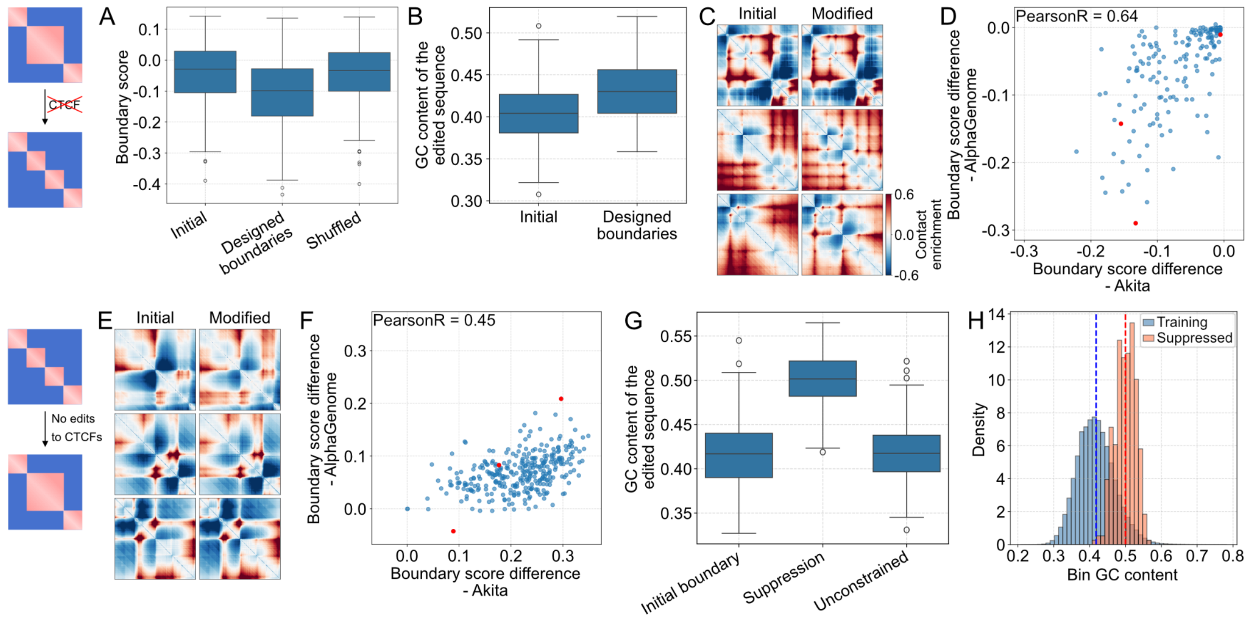


**Additional file 2: Figure S10. Boundary design without CTCF motifs and boundary suppression without modifying pre-existing CTCFs.**

**A)** Boundary scores for initial sequence predictions and for sequences edited to form boundaries in the absence of CTCF motifs, and for dinucleotide-preserving shuffled controls.

**B)** GC content within the edited subsequences for the initial sequences and after weak boundary design without introducing CTCF motifs.

**C)** Three examples of AlphaGenome predictions illustrating original sequences and their edited counterparts, which gain weak boundaries in the middle of the region despite lacking CTCF motifs. Examples are ordered by boundary score from weakest to strongest insulation (top to bottom, corresponding to red dots in (D)).

**D)** Quantitative validation of weak boundaries created without CTCF motifs. Boundary score differences (optimized – initial) computed using Akita and AlphaGenome predictions show concordance (Pearson's r > 0.6).

**E)** Representative AlphaGenome predicted contact maps for three sequences before and after locked-motif boundary suppression. Examples are ordered from strongest to weakest boundary suppression (top to bottom). The color scale is the same as in panel C.

**F)** Scatterplot of boundary score differences (optimized – initial) computed for Akita and AlphaGenome predictions. The cross-model correlation is moderate (Pearson's r > 0.4).

**G)** GC content within the edited region for initial sequences and after locked-motif boundary suppression. GC content increases significantly following suppression but not in unrestricted control.

**H)** Distribution of per-bin GC content across Akita training data (blue) and within the edited bins of boundary-suppressed sequences (orange), showing that optimized sequences remain within the training distribution.

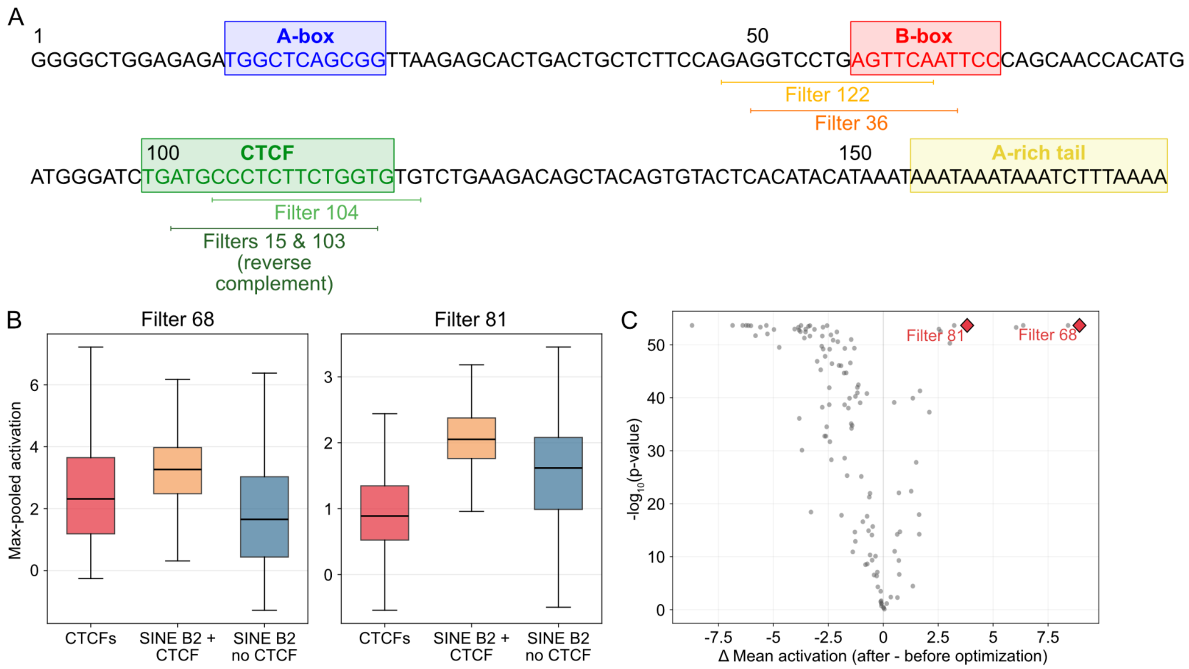


**Additional file 2: Figure S11. Akita recognizes SINE B2 elements and can use their features to suppress boundaries without deleting existing CTCF sites.**

**A)** Schematic of a B2_Mm2 SINE B2 element with annotated structural features: A-box, B-box, reverse-orientation CTCF motif, and A-rich tail. Filters strongly activated by SINE B2+CTCF elements were identified, and their learned sequence preferences were compared against the B2_Mm2 consensus sequence to assess correspondence (**Methods**). Identified matches include filters recognizing the B-box RNA Polymerase III internal promoter (filters 36 and 122) and the SINE B2-specific CTCF motif variant (filters 15, 103, and 104).

**B)** Activation of fourth convolutional layer filters (receptive field ~135 bp, sufficient to capture a SINE B2 element) across three sequence groups: isolated strong CTCF motifs, SINE B2 elements containing CTCF motifs (SINE B2+CTCF), and SINE B2 elements lacking CTCF motifs (SINE B2 no CTCF). Filters 68 and 81 show the highest specificity for the SINE B2+CTCF group.

**C)** Filters most specifically activated by SINE B2+CTCF elements relative to both SINE B2 elements lacking CTCFs and isolated CTCF motifs are also among the most strongly activated in boundary-suppressed sequences compared to their pre-optimization counterparts (filters 68 and 81, ranked 1st and 5th by activation change).

#### Additional file 3: Movie S1

**Additional file 3: Movie S1. Progressive transformation of genome folding during full sequence optimization.**

The movie begins with the predicted contact map for the initial DNA sequence. Below the map, the CTCF landscape is shown as forward-oriented motifs (black, positive bars) and reverse-oriented motifs (red, negative bars). Each subsequent frame displays the predicted contact map and corresponding CTCF landscape for the next sequence containing accepted edits. As optimization proceeds, the cumulative edits drive a progressive transformation of the predicted contact map toward the specified target folding pattern.
